# Amelioration of hemophilia B through CRISPR/Cas9 induced homology-independent targeted integration

**DOI:** 10.1101/2021.03.18.435908

**Authors:** Xi Chen, Xuran Niu, Yang Liu, Rui Zheng, Liren Wang, Lei Yang, Jian Lu, Shuming Yin, Yanjiao Shao, Yu Wei, Jiahao Pan, Ahmed Sayed, Xueyun Ma, Meizhen Liu, Fengxiang Jing, Mingyao Liu, Jiazhi Hu, Xiaohui Zhang, Dali Li

## Abstract

Site-specific integration of exogenous gene through genome editing is a promising strategy for gene therapy. However, homology-directed repair (HDR) only occurring in proliferating cells is inefficient especially *in vivo*. To investigate the efficacy of Cas9-induced homology-independent targeted integration (HITI) strategy for gene therapy, a rat hemophilia B model was generated and employed. Through HITI, a DNA sequence encoding the last exon of rat *Albumin* (*rAlb*) gene fused with a high-specific-activity Factor IX variant (R338L) using T2A, was inserted into the last intron of r*Alb* via recombinant adeno-associated viral (rAAV). The knock-in efficiency reached up to 3.66% determined by ddPCR. The clotting time was reduced to normal level 4 weeks after treatment, and the circulating FIX level was gradually increased up to 52% of normal during 9 months even after partial hepatectomy, demonstrating the amelioration of hemophilia. Through PEM-seq, no significant off-targeting effect was detected. Moreover, this study provides a promising therapeutic approach for hereditary diseases.

---

Gene therapy based on recombinant adeno-associated virus (rAAV) vectors provides a promising therapeutic approach. Recently, it has been shown promising treatment effects in clinical trials for variant diseases, such as inherited retinal dystrophy ^1^, spinal muscular atrophy^2^, hemophilia^3, 4^ and so on^5^. However, preclinical studies have shown that only shortterm effects have been observed when rAAV vector was delivered to target proliferating tissues in infant animal models^6, 7^. Moreover, a 3-year follow-up of clinical trial for hemophilia A showed a continuous decline of FVIII levels over time in adult patients^8^, indicating that rAAV-mediated gene replacement therapy may exist problem with the durability of treatment, because rAAV vectors predominantly form episomes which would lose during cell proliferation^9^.

CRISPR/Cas9 system is a widely used genome editing technology which induces sitespecific double-stranded break (DSB) with the guidance of single-guide RNA (sgRNA). DSB is repaired either through an error-prone nonhomologous end joining (NHEJ) pathway or a precise HDR pathway in the presence of exogenous DNA templates^10^. CRISPR/Cas9-mediated HDR strategy has been successfully used to ameliorate genetic disorders through correction of disease-causing genetic mutations or targeted integration of exogenous DNA at a target site, including hemophilia^11, 12, 13^, phenylketonuria^14^ and ornithine transcarbamylase deficiency (OTCD)^15, 16^. However, although HDR is highly accurate, its efficiency is limited even delivered through rAAV vector whose single-stranded DNA genome is considered to induce a higher HDR rate^17^.

Since NHEJ is active throughout the cell cycle^18^, NHEJ-mediated targeted integration strategy provides an alternative approach to insert exogenous DNA in adults. In addition, this strategy is independent of homologous arm to insert relatively larger DNA fragments especially using an rAAV vector whose maxima capacity is about 4.7kb (≤5 kb)^19^. Recently, a CRISPR/Cas9-mediated homology independent targeted integration (HITI) strategy was developed to efficiently targeted knock-in exogenous genes in dividing and non-dividing cells. Importantly, through an innovative design of insertion of CRISPR/Cas9 recognition sequences within donor templates, the preferred orientation of inserted DNA could be determined with a high probability through HITI. Using this strategy, retinitis pigmentosa retina was partially rescued in a rat model, suggesting that HITI is also functional in nondividing cells *in vivo*^20^.

To investigate the potential of HITI to treat diseases in target organs besides retina, we generated a hemophilia B (HB) model via CRISPR/Cas9 edited *F9* gene in rats, since Factor IX (F□) expresses mainly in hepatocyte which is a very critical cell type for gene therapy not only to treat diseases caused by hepatocyte dysfunction but also for the production of therapeutic secreting factors. Hemophilia B is an ideal candidate to test novel strategies for gene therapy targeting hepatocytes, as our previous study has demonstrated that moderate correction efficiency (>0.56%) would significantly ameliorate disease symptoms in mice^11^. Moreover, F□ is a secreting protein which is a typical factor representing the enzyme replacement therapy. In this study, we tested the feasibility of HITI strategy to treat hemophilia B through targeted insertion of a high-specific-activity F□ variant (*F9* Padua, R338L) into the last intron of rat *Alb* (r*Alb*) gene locus. Through rAAV8 delivered Cas9/sgRNA and template in *F9* deficient adult animals, the circulating FIX level was increased up to 52% of normal level during 9 months observation, demonstrating the amelioration of hemophilia B.

## Results

### Generation of *F9^Δ130/Δ130^* rat strains

The previously reported mouse model of hemophilia B mimics thrombin dysfunction well, but does not show a phenotype similar to human spontaneous bleeding^11^. Compared with mice, rats are larger in size and their physiology is closer to that of humans^21^. Since the rat hemophilia B model has not been reported, we first generated a rat HB model through injection of Cas9 mRNA and two sgRNAs targeting exon 2 and intron 2 of *F9* gene respectively into zygotes (Fig. 1a). The sgRNA sequences and primers are shown in Supplementary Table 1. Sanger sequencing results of the genomic DNA (gDNA) validated the successful generation of *F9* KO rats with 130bp deletion which produced a premature stop codon at the end of exon 2 of r*F9* gene (hereafter called the strain as *F9^Δ130/Δ130^*) (Fig. 1b). Compared with wild-type (WT) rats (n=5), *F9^Δ130/Δ130^* rats did not detect any *F9* mRNA expression in liver by RT-PCR (Fig. 1c). The average activated partial thromboplastin time (aPTT) of 8-week-old WT rats was 20.12 ± 1.217s, which was significantly prolonged to 68.04 ± 5.739s in *F9^Δ130/Δ130^* rats. No significant difference in average prothrombin time (PT) was observed between these two groups (Fig. 1d). These rats were subjected to a tail-clip challenge for further confirmation of the coagulation function. The bleeding volumes of *F9^Δ130/Δ130^* rats within 6 minutes after the tail-clip were more than 2.8 times over that of WT rats (Fig. 1e). All *F9^Δ130/Δ130^* rats died within 48 hours after tail-clip without treatment, but no death was observed in WT rats. In addition, intra-articular bleeding and swelling were observed in 20% of *F9^Δ130/Δ130^* rats within 2 to 3 months of age (Fig. 1f). It is in line with the symptom of human hemophilia arthritis which has not been observed in HB mice^22^. These results indicate the successful generation of HB rat model which is more closely recapitulating human HB symptoms than mouse model.

**Fig.1.**
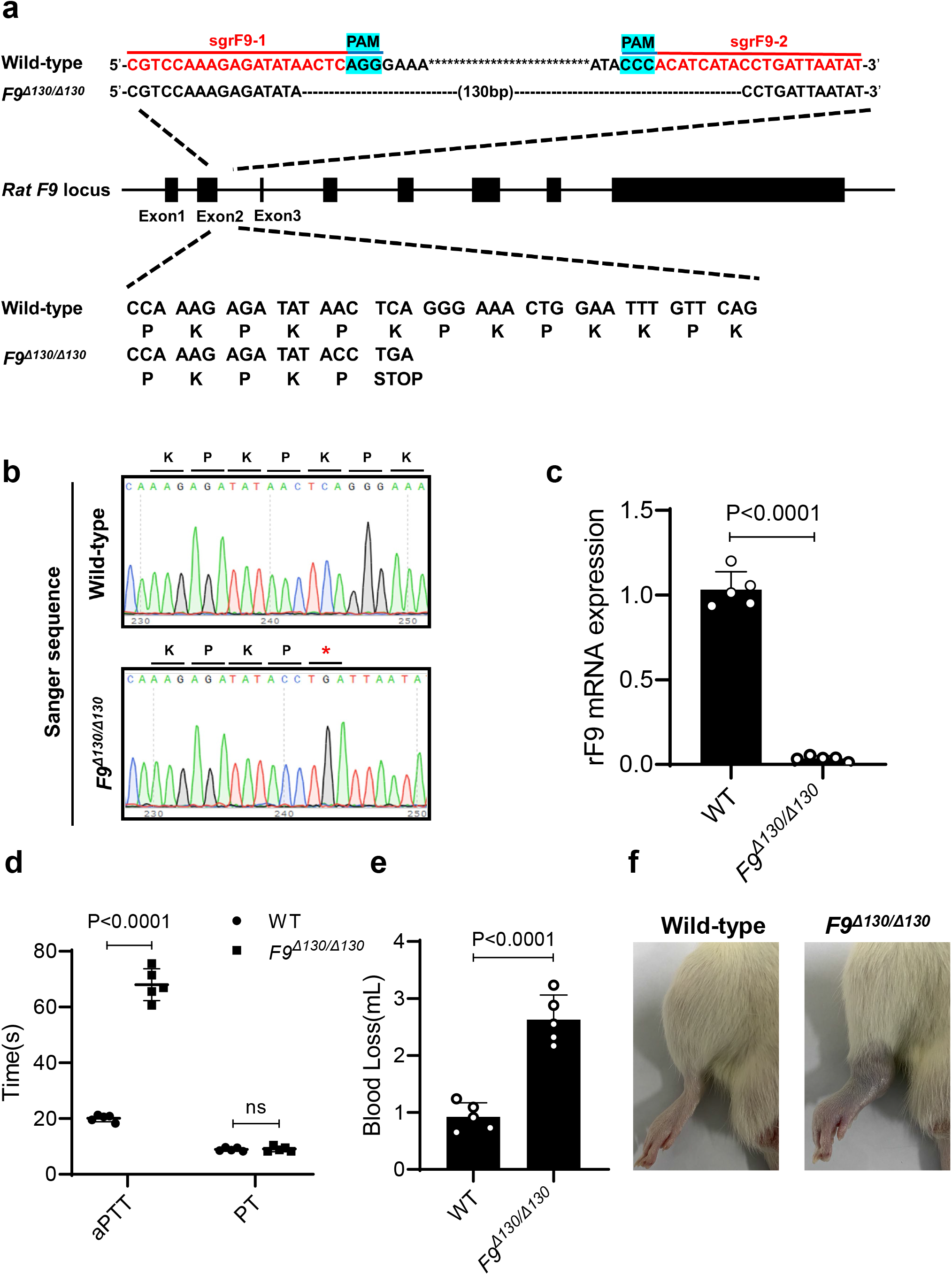
Generation of hemophilia B rats. **a** Schematic diagram of *F9* gene knock-out mediated by CRISPR/Cas9 in SD rats shows the design of target sites (sgRNA sequences are indicated by the red line and the PAM sequence is labeled in blue) and the mutant sites of *F9^Δ130/Δ130^* rat strains are represented with dash lines. The reading frames of the WT and mutant *F9* sequences are shown and the 130-bp deletion in *F9* gene results in an early STOP codon. **b** Sanger sequencing of the targeted *F9* sites of WT and *F9^Δ130/Δ130^* rats. Asterisk in red represents stop codon. **c** Expression of *F9* mRNA in the liver tissues of wild type (WT) and *F9^Δ130/Δ130^* rats (n=5) by RT-PCR. **d** Blood coagulation assessed by activated partial thromboplastin time (aPTT) and prothrombin time (PT) were measured in WT and *F9^Δ130/Δ130^* rats (n=5). **e** Measurement of bleeding volume after tail clipping in WT and *F9^Δ130/Δ130^* rats at 8 weeks of age (n=5). **f** Hemarthrosis caused by spontaneous bleeding in *F9^Δ130/Δ130^* rats. Representative photographs of WT (left) and *F9^Δ130/Δ130^* rats (right) are shown. Intra-articular bleeding and swelling were observed in 20% of *F9^Δ130/Δ130^* rats within 2 to 3 months of age. (Values and error bars reflect the means and SD. of five independent experiments. P value was determined by two-tailed Student’s t-test. ns, no significant difference by two-tailed Student’s t test.)

### Construction of CRISPR/Cas9-mediated rAAV-HITI vector

Since Albumin is the most abundant protein secreted by hepatocytes, we decided to insert a hyperactive human FIX-R338L mutation (also called h*F9* Padua variant, h*F9*p) cDNA^23^ into the *Alb* locus. Some previous reports have shown *Alb* locus is suitable, but their strategy disrupted endogenous *Alb* expression when the alleles were inserted by gene of interest^13, 24^ As the last exon of *Alb* is very short (39bp), we decided to insert a DNA fragment following a splicing acceptor sequence through HITI strategy. The donor template was constructed in order as this: a splice acceptor (SA) sequence, r*Alb* exon 14, T2A sequence encoding selfcleaving peptide, h*F9*p cDNA and bovine growth hormone (bGH)-polyA sequence. More importantly, the donor template was flanked by two homonymous Cas9/sgRNA recognition sequences which were reversely orientated to endogenous Cas9/sgRNA recognition sequence in r*Alb* intron (Fig. 2a). This design is very important to increase the chance of insertion with correct orientation^20^. As shown in Supplementary Fig. 1, the principle of HITI strategy is to use the Cas9/sgRNA to simultaneously cleave the target sites of genome and donor vector, and the linearized donor sequence inserted in the desired orientation prevented the secondary cleavage of Cas9 due to the damage of target sites, while the insertion of donor DNA in the undesired orientation would reassemble the complete target sites, which lead to the secondary cleavage to remove the inserted sequence and enhance the chance for desired donor DNA insertion. Therefore, HITI mainly inserts the foreign DNA sequences in the desired orientation.

**Fig.2.**
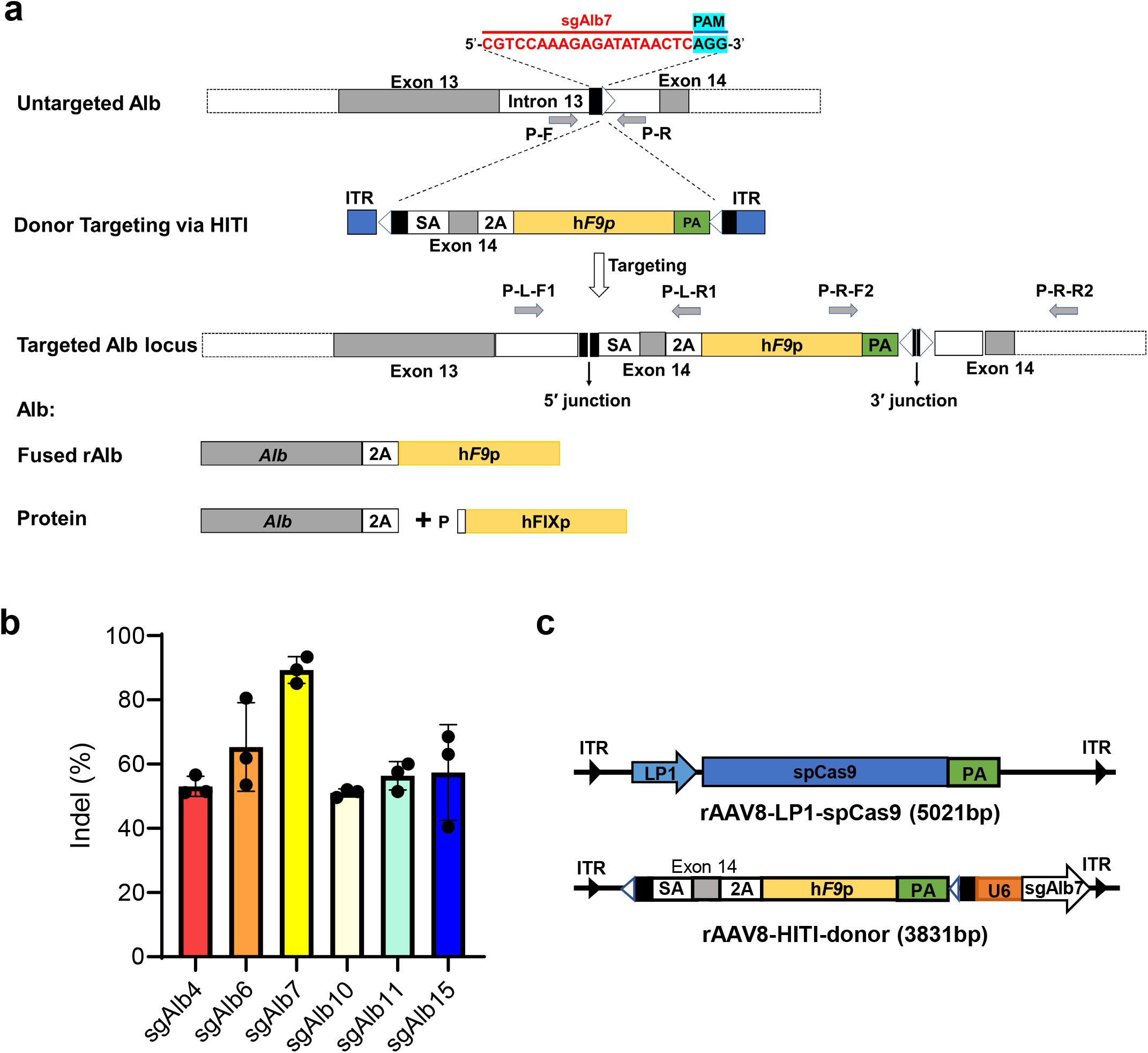
Vector design and genome editing of r*Alb* locus via HITI. **a** Schematic illustration of the site-specific integration in the r*Alb* intron 13 using CRISPR/Cas9-mediated HITI. The double-cut donor vector (rAAV8-HITI-donor) which contains a single-guide RNA with PAM sequences tagged close to the 5’ ITR, a splice acceptor (SA) sequence, a T2A sequence, r*Alb* exon 14 (39bp) located between T2A sequence and SA sequence, a codon-optimized h*F9* DNA (CDS) sequence that carries a hyperactive Padua mutation (R338L) and the bovine growth hormone polyA (bGH-polyA) flanked by T2A sequence and a same-orientation guide RNA recognition sites with PAM sequences, and a U6-sgAlb7 cassette placed close to the 3’ ITR. The fused mRNA of r*Alb* and h*F9* genes were co-expressed by r*Alb* promoter and the fusion protein is split by the self-cleavage of T2A peptide. 5’ and 3’ junctions indicate the splicing sites of donor sequence at r*Alb* locus. Blake pentagon, Cas9/gRNA target sequence. **b** The frequencies of CRISPR/Cas9-induced indels at 6 sgRNA sites were analyzed by Synthego website. Error bars represent means ±SD, n=3 for each group. **c** Schematic representation of rAAV plasmids used for treatment *in vivo*. The rAAV8-LP1-spCas9 expressing humanized spCas9 by using a liver-specific promoter (LP1). rAAV8-HITI-donor, the double-cut donor template and sgRNA targeting the r*Alb* intron 13 was inserted into an rAAV vector.

To screen for highly efficient sgRNAs within last intron of r*Alb* gene, six sgRNAs predicted by Benchling’s website (www.benchling.com) were tested in rat pheochromocytoma PC12 cells (Supplementary Fig. 2a). Though both T7 endonuclease I (T7EI) assay and ICE analysis (ice.synthego.com/#/) of Sanger sequencing chromatograms of the PCR products, we chose the most efficient sgAlb7 with 89.3% of indels for further studies (Fig. 2b and Supplementary Fig. 2b, c). According to the design of HITI, we next constructed a double-cut donor plasmid containing the donor template (hereafter called HITI-donor) and a pSpCas9(BB)-2A-GFP (PX458) expressing the Cas9 protein and sgAlb7, respectively (Supplementary Fig. 3a). All plasmid constructs were verified by sequencing.

### Evaluation of HITI-mediated h*F9*p integration in PC12 cells

We first evaluate the feasibility of HITI-mediated h*F9*p cDNA targeting in the r*Alb* intron 13 by transiently transfecting rat PC12 cells with PX458 and HITI-donor plasmids *in vitro* following FACS sorting of GFP positive cells (Fig. 2c), which induced an average indel rate of 27.3% by tagged sequencing and achieved a high targeted integration efficiency (□ 17.68%) in the directed orientation as determined by droplet digital PCR (ddPCR, Supplementary Fig. 3b and Supplementary Fig. 9a). We further used tagged sequencing to verify the fidelity of HITI *in vitro*. The result showed that the majority of forward-directed splicing sequences at 5’ and 3’ junction sites were precise (81.35% for 5’ junction site and 67.98% for 3’ junction site, respectively). Only a small amount of imprecise integration was observed, including fragment deletions, insertions, and point mutations. However, this imprecise splicing at target site was mainly located in the non-coding region of the donor sequence, which thus did not disrupt the coding of r*Alb* and h*F9*p proteins (Supplementary Fig. 2c). The above results indicate that HITI strategy can efficiently and precisely target h*F9*p cDNA into intron 13 of r*Alb* gene *in vitro*.

### HITI-mediated high levels of hFIX protein via rAAV8 delivery to ameliorate hemophilia B *in vivo*

Encouraged by the above results, we further explored the feasibility of this strategy for the treatment of hemophilia B *in vivo*. As our previous study showed that AAV8 transduced efficiently in neonatal rat hepatocytes ^25^, we also confirmed in this study that 1×10^11^ genome copies (gc) of rAAV8-CMV-EGFP could infect and express very well in 8-week-old rats via tail vein injection (Supplementary Fig. 4). Then, we generated a dual rAAV8s system: a vector coding *Streptococcus pyogenes* Cas9 (spCas9) protein with the LP1 promoter (hereafter called rAAV8-LP1-spCas9)^25^ and another containing the double-cut template and U6-sgAlb7 expression cassette (hereafter called rAAV8-HITI-donor) (Fig. 2c). Five different groups had been set up: in low-HITI group, *F9^Δ130/Δ130^* rats received 2E +11 gc/rat rAAV8-LP1-spCas9 and 4E +11 gc/rat rAAV8-HITI-donor; high-HITI group, *F9^Δ130/Δ130^* rats received 2E +12 gc/rat GC rAAV8-LP1-spCas9 and 4E +12 gc/rat GC rAAV8-HITI-donor; donor group, *F9^Δ130/Δ130^* rats received 4E +12 gc/rat rAAV8-HITI-donor; untreated group and WT group made up of *F9^Δ130/Δ130^* rats and WT rats receiving 800 μL PBS only, respectively.

After rAAV injection, we collected plasma to measure the activity and levels of hFIX protein. Activated partial thromboplastin time (aPTT) assay was employed to monitor the function of plasma hFIX protein. Four weeks after rAAV injection, the APPTs of low-HITI group and high-HITI group were reduced to 35.6 ± 8.5s (n=3) and 25.9 ± 4.4s (n=3) respectively, which were significantly shorter than those of the donor group (63.28 ± 0.725 s, n=3) and the untreated group (66.02 ± 5.111 s, n=5), and tended to that of WT group (23.50 ± 1.208 s, n=5) from 12 weeks after the injection until the end (Fig. 3b).

**Fig.3.**
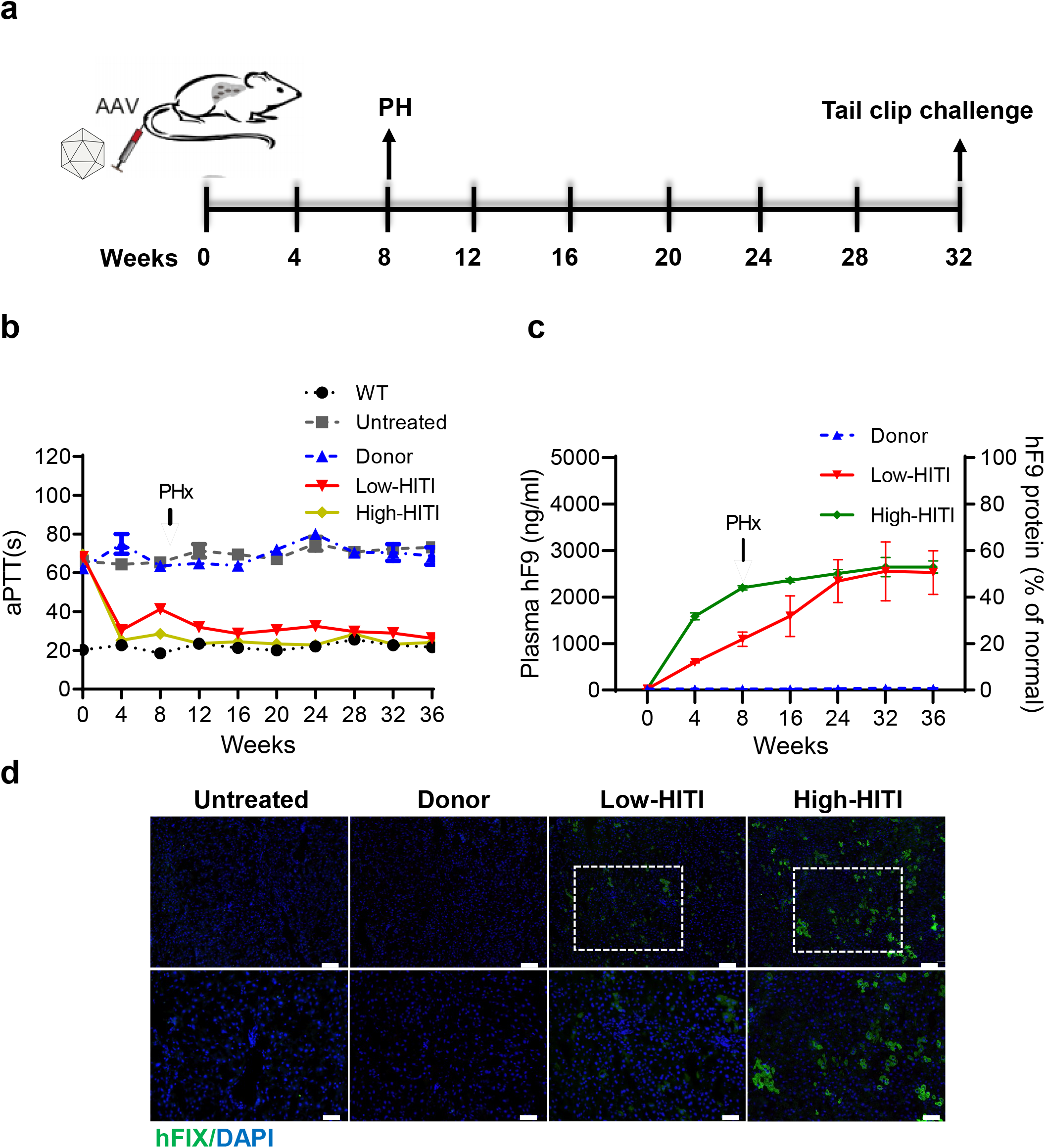
Efficient, sustained expression and activity of hFIX protein in HITI-treated rats. **a** Schematic diagram of the experimental design for rAAV injection and analysis. PHx, partial hepatectomy. **b**, **c** hFIX protein activity and levels in the plasma were assayed by aPTT and ELISA, respectively. The measurements were performed from 4 to 36 weeks after rAAV injection with subsequent PHx. hFIX levels were shown as the percentage of normal hFIX levels in the plasma. Error bars represent means ±SD. WT group, n=5; untreated group, n=5; donor group, n=3; low-HITI group, n=3; high-HITI group, n=3. **d** Immunofluorescence staining with antibodies against hFIX (green) and with DAPI (blue) on rat liver sections collected from PHx 8 weeks after rAAV injection. Scale bars, 200 μm (top), 100 μm (bottom).

To further investigate the actual hFIX protein level in treated rats, enzyme-linked immunosorbent assay (ELISA) was leveraged. The plasma hFIX level of both the low-HITI and high-HITI groups increased dramatically from 4-week and reached plateau at 32-week and kept the high level to the end of the experiments (Fig. 3c). Interestingly, both the low-HITI and high-HITI groups reached over 50% (51%, 2553 ± 632.4 ng/mL in low dose group; 53%, 2647 ± 205.5 ng/mL in high dose group), although the high-HITI group exhibited a sharper increase rate (Fig. 3c). As expected, the donor and untreated groups, plasma hFIX levels were barely detectable (~20 ng/ml). Additionally, partial hepatectomy (PHx) showed no impact on the stable activities and evaluated levels of hFIX protein in the plasma of all treated rats, indicating stable gene integration (Fig. 3b, c). Liver samples from PHx of all groups were analyzed for hFIX expression by immunofluorescence (Fig. 3d) and immunohistochemistry (Supplementary Fig. 5a). These results showed that significant hFIX-positive hepatocytes were presented in both low dose and high dose treated HB rats in a dosedependent manner which was further confirmed by western blotting via hFIX antibody (Supplementary Fig. 5b, c). At 36 weeks after treatment, a tail-clip challenge was performed to further confirm the therapeutic effect. Similar to WT group (n=6), all treated rats (n=6) lost 3-fold fewer blood volumes than rats from the untreated group (n=5) (Supplementary Fig. 6a). All rats from the untreated group (5 out of 5) and the donor group (3 out of 3) died within 48 hours after tail clip, while no deaths occurred in all treated rats and WT rats (Supplementary Fig. 6b). These results demonstrate that HITI strategy could effectively and persistently restore the expression and activity of hFIX protein, and significantly ameliorate the coagulation function in adult HB rats.

### Genomic analysis of rAAV-based hF9p cDNA knock-in at rAlb intron 13 via HITI *in vivo*

To evaluate the cleavage activity of CRISPR/Cas9 *in vivo,* we analyzed on-target indel frequency in gDNA of liver tissue from treated rats 8 weeks after rAAV injection via tagged sequencing. The indel frequencies were 3.2%-5.3% in low-HITI group and 10.3%-16.7% in high-HITI group respectively, whereas no indels were detected in untreated and donor groups (Fig. 4a). In addition, we harvested the gDNAs of other 4 organs except for the livers from untreated and high-HITI treated rats 36 weeks after rAAV injection and did not observe any detectable indels (Supplementary Fig. 7). The results demonstrated that Cas9/sgAlb7 delivered by rAAV8 can induce efficient and tissue-specific cleavage at the target site *in vivo*. To further detect the HITI-mediated targeted integration at the target site, we designed 2 pairs of primers (P-L-F/R and P-R-F/R, Supplementary Table 2) to amplify the 5’ and 3’ junctions by PCR (Fig. 2a). Gel electrophoresis showed the correct integration bands (5’ junction: 529bp and 3’ junction: 1300bp) in livers gDNAs of all treated rats (Supplementary Fig. 8a). Furthermore, we performed ddPCR to quantitate HITI-mediated targeted integration efficiency 8 weeks after rAAV injection (Supplementary Fig. 9). The efficiencies of low-HITI group and high-HITI group were 0.52%-1.14% and 1.31%-3.66%, respectively (Fig. 4a).

**Fig.4.**
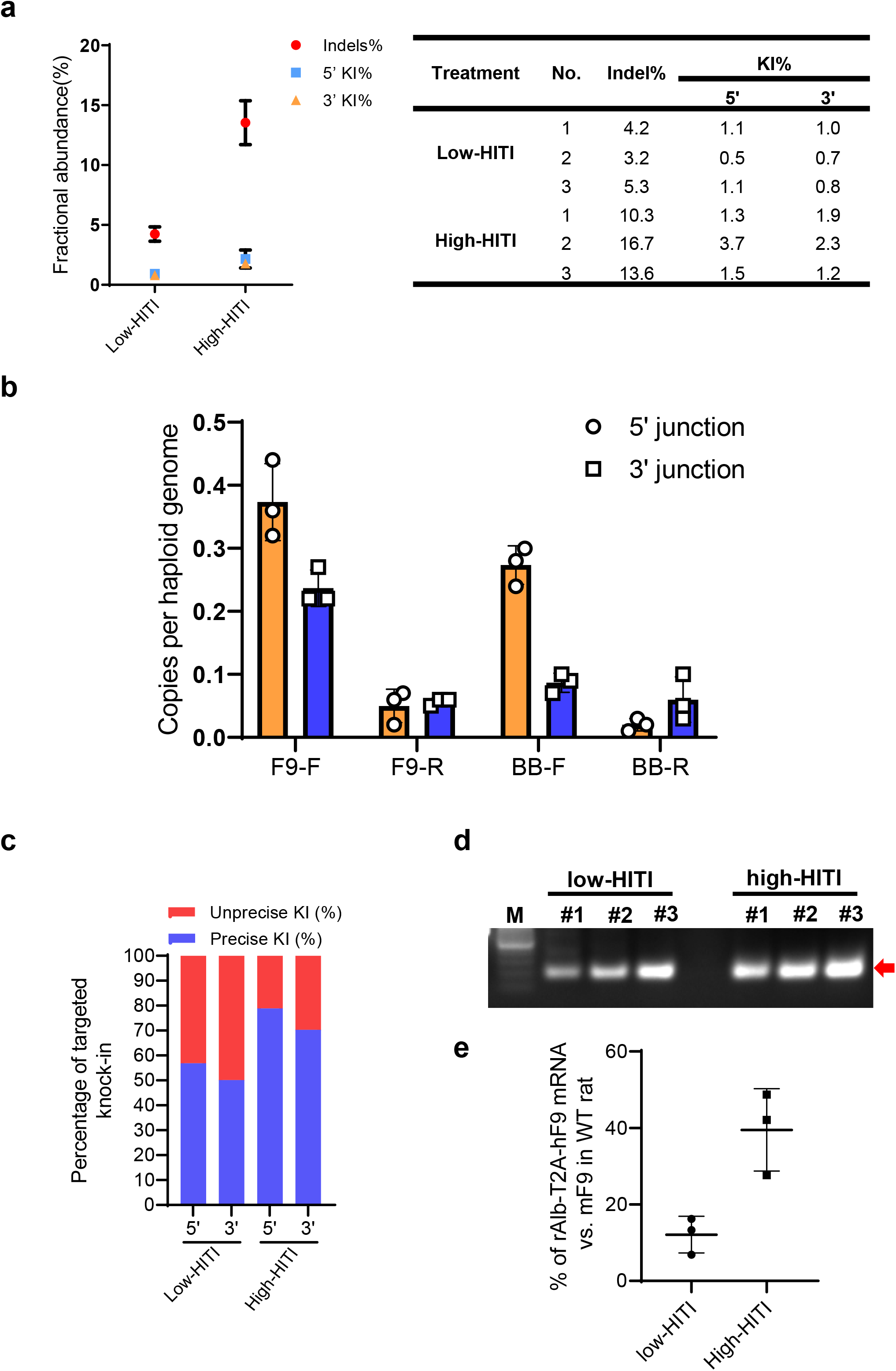
Indel and HITI-mediated gene integration efficiency and specificity of *hF9* expression. **a** The indel frequency of sgAlb7 and h*F9*p transgene knock-in efficiency (KI%) at the targeted *Alb* locus as determined by tagged sequencing and ddPCR analysis, respectively. Error bars represent SD from n = 3 replicates. **b** Quantitation of copy numbers for integration types of h*F9*p cDNA and rAAV backbone in forward and reverse orientation in r*Alb* target site by PEM-seq. F9-F, insertion of h*F9*p cDNA in the forward orientation; F9-R, insertion of h*F9*p cDNA in the reverse orientation; BB-F, insertion of rAAV backbone in the forward orientation; BB-R, insertion of rAAV backbone in the reverse orientation; Error bars represent SD, n=3 for HITI-treated group. **c** Frequency of the fidelity at 5’ and 3’ junction sites of h*F9*p knock-in by HITI. **d** PCR analysis showing a successful fusion of r*Alb* and h*F9*p mRNAs 8 weeks after rAAV injection. The cDNA produced by reverse transcription was served as a template for amplification of the fusion transcript of r*Alb*-T2A-h*F9*p. **e** Quantitative RT–PCR of liver total RNA from HITI-treated rats. *rAlb-T2A-hF9* Pauda mRNA levels are relative to endogenous m*F9* mRNA levels of WT rats. RNA samples were analyzed in duplicate.

To comprehensively map all possible genome-editing outcomes with an unbiased approach, we applied a high-throughput method called primer-extension-mediated sequencing (PEM-seq)^26^. About 20 μg fragmentated liver gDNA was used for each PEM-seq library, and the bait primer was placed within 200 bp from the target site (Supplementary Fig. 10a; see Materials and methods for details). Three independent biological replicates for each treatment were generated and combined for translocation junctional hotspots analysis. We analyzed the mapped reads and defined the ratio of reads containing the expected h*F9*p sequence and total mapped reads as HITI-mediated targeted integration efficiency. Compared with ddPCR, PEM-seq data showed relatively lower efficiency with 0.43%-0.64% (low-HITI group) and 0.94%-1.66% (high-HITI group) (Supplementary Fig. 10b). Additionally, a more detailed analysis of PEM-seq data showed different targeted integration events at the target site, including the forward and reverse insertions of h*F9*p cDNA and rAAV backbone. Among them, the insertion efficiency of h*F9*p cDNA in the desired orientation is 7.5-fold higher than that in the reverse orientation (Fig. 4b), which is consistent with a previous report^20^. Additionally, we also evaluated the fidelity of HITI-mediated targeted integration *in vivo* via PEM-seq. Consistent with the result *in vitro,* the majority of the integration of h*F9*p cDNA was the seamless integration with very high ratio of 56.86%-78.87% at 5’ junction site and 50.10%-70.28% at 3’ junction site (Fig. 4c and Supplementary Fig. 8b). To further confirm whether HITI-mediated targeted integration affects the correct splicing of r*Alb* and h*F9*p gene, we extracted mRNA of livers from all groups 8 weeks after rAAV injection and performed RT-PCR to detect the r*Alb*-T2A-h*F9*p fusion transcriptional products. Consistent with the results of PEM-seq, gel electrophoresis result showed the correct bands indicating the donor integration in all treated rats (Fig. 4d). The levels of the r*Alb*-T2A-h*F9*p fusion mRNA in all treated rats were approximately 4.4%-33.3% of r*F9* mRNA levels present in WT rats (Fig. 4e). Sanger sequencing further verified the correct splicing of r*Alb* exon13 and exon14-T2A-h*F9*p (Supplementary Fig. 11). The above results showed that h*F9*p cDNA could be efficiently and precisely inserted into the r*Alb* intron 13 through rAAV-delivered HITI strategy *in vivo,* which generated the correct fusion transcripts.

### Safety evaluation of genome editing therapy for hemophilia B via rAAV-CRISPR

CRISPR/Cas9-induced off-target activity is a major safety concern for gene editing therapy *in vivo*. To examine the *in vivo* off-target effects of the CRISPR/Cas9 system, we selected 11 predicted high-activity off-target sites of sgAlb7 using Benchling’s website (Supplementary Table 3). 8 weeks after rAAV injection, we amplified the liver gDNAs from the untreated and high-HITI groups. Tagged Sequencing of the top 11 predicted off-target sites showed no significant increase in off-target cutting in all treated rats compared to the untreated rats (Fig. 5a and Supplementary Table 4). To further detect the off-target effects at the genome-wide level, we used PEM-seq to capture the off-target sites of sgAlb7. Except for the translocation between the target site and donor sequence, we did not observe translocation at any off-target site including the predicted ones (Fig. 5b). These results suggested that sgAlb7 possessed a high specificity for its expected target site. Previous reports have suggested a risk of immune response and/or genomic toxicity induced by rAAV vector^27^. Hence, we evaluated liver toxicity of rAAV-mediated gene therapy in hemophilia B rats. Further studies demonstrated no significant difference in plasma aspartate transaminase (AST), alanine transaminase (ALT) and Albumin (ALB) levels (Fig. 5b) and the transcription of inflammatory factors (Fig. 5c) between the treated and untreated rats, suggesting the good tolerance of rAAV vectors, which was confirmed by histology analysis 8 weeks after rAAV injection (Fig. 5d). In conclusion, the HITI strategy delivered by rAAV did not induce severe off-target effects and liver toxicity in hemophilia B rats.

**Fig.5.**
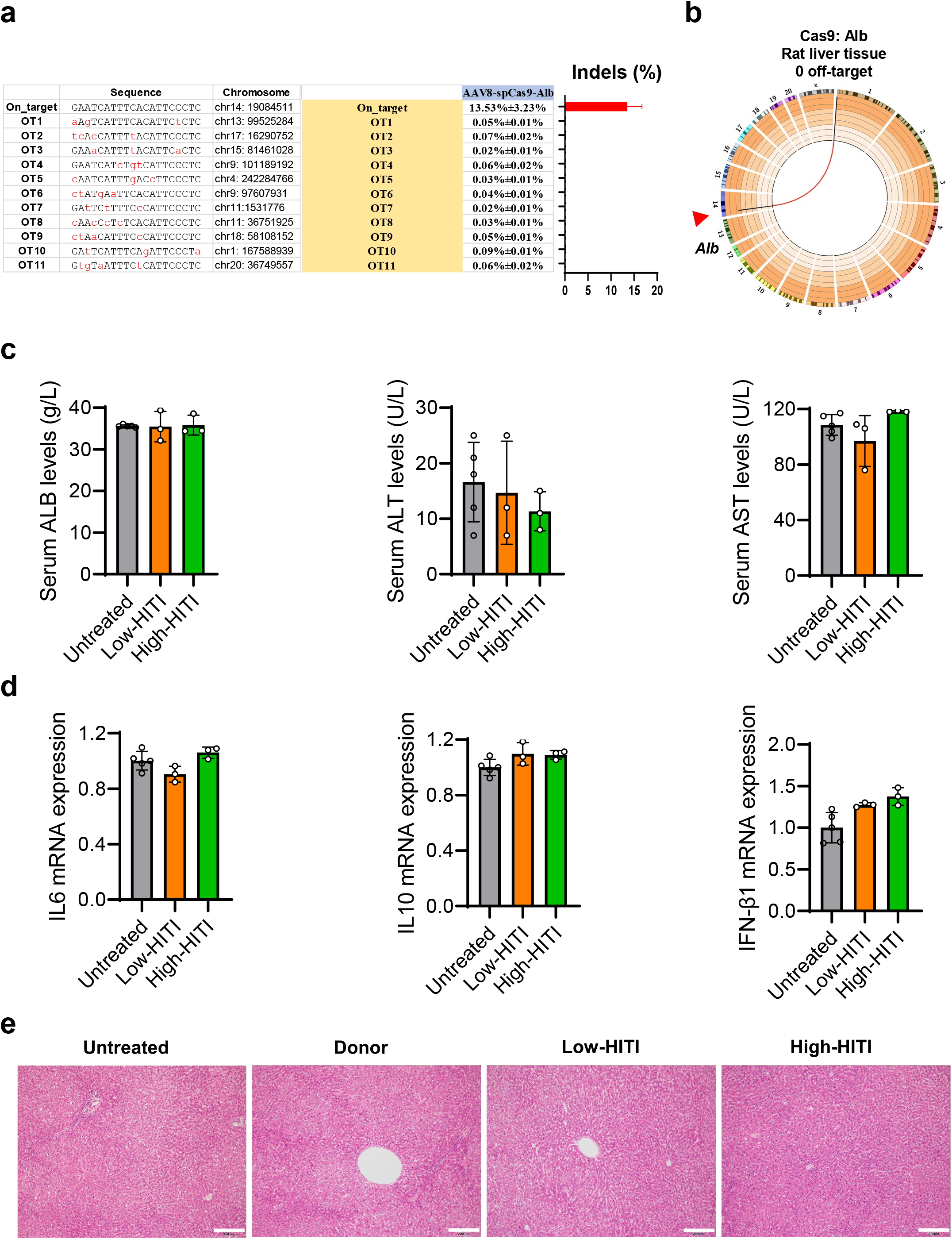
Safety evaluation of gene targeting therapy for hemophilia B via rAAV8.HITI. **a** Off-target analysis by tagged sequencing in liver gDNAs. A list of on- and off-target analysis using gDNA isolated from the rat livers by tagged sequencing. The nucleotide letters shown in red are the individual mismatches in predicted off-target sites. **b** Universal bait detection of off-targets for sgAlb7. Circos plot (circos.ca) on custom log scale showing the genome-wide profile of Cas9: r*Alb* junctions. The bin size is 5 Mb, and 8,751 unique junctions are shown from two independent libraries. Chromosomes were showed with centromere to telomere in a clockwise orientation. Red arrow indicated the Cas9: r*Alb* cleavage site. For the accuracy of PEM-seq analysis, we artificially placed the complete donor DNA sequence into the front of chromosome 1 of *Alb* as reference. Colored lines showed the hotspot between the genomic on-target site and the donor DNA sequence. **c** Quantification of spCas9 vector genome in the liver by qPCR. Error bars represent SD, n=6.Serum markers of liver toxicity, such as total ALB, ALT and AST levels were determined 32 weeks after rAAV injection. No significant differences were observed among all groups. Error bars represent means ±SD. untreated group, n=5; low-HITI group, n=3; high-HITI group, n=3. **d** mRNA levels of inflammatory cytokines from livers of HITI-treated rats were determined by RT-PCR. Error bars represent means ±SD. untreated group, n=5; low-HITI group, n=3; high-HITI group, n=3. **e** Histological analysis by hematoxylin and eosin (HE) staining of liver sections from untreated group (n=5), donor group (n=5), low-HITI group (n=3) and high-HITI (n=3) group 8 weeks after rAAV injection. The images represent the liver section of one rat from each group. Liver tissues showed no histological abnormalities in either HITI-treated groups as compared to untreated and donor groups’ rats.

## Discussion

Site-specific integration of therapeutic gene provides an ideal gene therapy method for patients with mutations elsewhere in disease-causing gene. However, *in vivo* HDR-mediated targeted integration is still infeasible due to low editing efficiency, especially in adults. To overcome the limit, the NHEJ-based HITI strategy has been proven to be capable of targeted integration of exogenous genes in desired orientation in both dividing and non-dividing cells. In this work, we generated a novel hemophilia B rat model (*F9^Δ130/Δ130^*) with spontaneous bleeding and hemophilia arthritis. Using HITI strategy via rAAV delivery, we successfully restored hemostasis in the adult rats with hemophilia B, suggesting HITI-mediated knock-in of therapeutic gene at r*Alb* intron 13 might be a favorable gene therapy strategy for various inherited metabolic disorders.

The establishment of disease animal models similar to the pathological features of human genetic disorders is of great significance in the assessment of clinical effectiveness and safety. Previous hemophilia gene therapy studies mainly used the mouse models^11, 24, 28, 29^, which were difficult to collect sufficient blood and tissue samples for analysis. Rats are at least 10 times larger than mice, and repeated blood sampling is possible, which can thus set up a smaller number of animals^30^. In physiological and pathological terms, rats are more analogous to humans than mice^31^. Besides severe coagulation dysfunction, our *F9^Δ130/Δ130^* rat model showed the symptom of intra-articular bleeding and swelling similar to human patients two months after birth, which was not reported in HB mouse model. Therefore, the *F9^Δ130/Δ130^* rat strain is a perfect model to imitate human disease.

Recent studies have successfully used site-specific targeted integration strategies to restore hemostasis in HB mouse models^9^. Site-specific integration of *hF9* cDNA has been achieved through nuclease dependent or independent strategies, such as rAAV donor mediated HDR, ZFN or CRISPR/Cas9 stimulated HDR-mediated insertion to ameliorate hemophilia B in mouse models ^9, 13, 32^. However, these studies showed a lower HDR efficiency in the adult liver compared with that in newborns. Recently NHEJ-based target integration strategies have been reported, such as ObliGate^33^, PITCh^34^, HMEJ^35^, HITI^20^ and SATI^36^, which provide alternative tools for the targeted integration of exogenous DNA in nondividing cells. However, the design of ObliGate is complex; PITCH cannot preclude NHEJ-mediated integration in a random direction; HMEJ exists the repeated insertion of homologous arm mediated by NHEJ; SATI showed less efficient in the liver (~2%). Compared with these strategies, NHEJ-based HITI is a more precise knock-in strategy in a desired orientation in non-dividing cells. In our study, we used HITI strategy to target h*F9* cDNA to r*Alb* intron 13 in an adult rat HB model, since *Alb* locus exhibited very high transcriptional activity. To minimize the impact of the ALB expression, we added *Alb* exon 14 (only 39bp) to the N-terminal of h*F9*p cDNA, which achieved the co-expression of r*Alb* and h*F9*p through T2A fusion. As a result, we observed the correct fusion transcript of r*Alb*-T2A-h*F9*p and no significant change in serum ALB levels.

So far, there is no effective quantitative method to detect the targeted integration efficiency of large fragments. Traditional PCR is difficult to quantitate targeted integration efficiency because unbiased amplification cannot be achieved. Recently, based on linear-amplification mediated PCR (LAM-PCR), some groups developed novel methods to measure HDR-mediated targeted integration efficiency^13, 29^, such as ligation-mediated PCR (LMU-PCR). However, the strategies apply restriction enzymes to digest DNA, which is not applicable to analyze the editing events without the restriction enzyme sites, such as translocation between target site and genome-wide or rAAV backbones, the reverse integration. Hence, the integration efficiency is possible to be over-estimated. ddPCR is another highly sensitive assay for the quantification of DNA and RNA targets, which has been used to quantitate NHEJ-targeted integration efficiency *in vivo*^37^. However, individual assays are required for each integration event, and multiplexing can be difficult. PEM-seq is a high-throughput sequencing method to comprehensively evaluate the editing events of genome editing^26^.

By employing a large amount of sonicated DNA (over 20 ug for each sample), one-round of linear amplification, and ligation with unique molecular barcode, PEM-seq can precisely quantify almost all the translocation events on target site, simultaneously, including on-target and off-target events. In our study, the PEM-seq assay achieved a lower targeted integration efficiency than ddPCR, which may be caused by limited copy number assays and missing other translocation events of ddPCR including the insertion events of h*F9*p cDNA in reverse orientation and rAAV backbone. In addition, a recent study ^37^ reported that NHEJ-mediated targeted integration of *F8* BDD cDNA at the *Alb* locus resulted in the forward and/or reverse insertion of *F8* BDD cDNA and rAAV backbone with equal efficiency at the target site. However, our PEM-seq analysis showed that the integration efficiency of the donor sequences in the desired orientation was 7.5-fold higher than that in the reverse orientation (Fig. 4b), indicating that HITI can further improve the forward insertion efficiency of donor sequences compared to NHEJ. Moreover, besides the sequences corresponding to parental genomic DNA and HITI-mediated insertion of h*F9*p cDNAs, partial rAAV backbone sequences were found to be inserted into the target sites. Given that rAAV ITRs promote vector integration at DSBs^38^, we also observed parts of ITR-flanking h*F9*p cDNAs inserted into the target sites (Supplementary Fig. 10c). Hence, we speculated that part of the therapeutic effects might be attributed to the integration of rAAV vector bearing T2A-h*F9*p cDNAs. These results indicate that PEM-seq can comprehensively evaluate genome editing effects such as on- or off-target insertion efficiency, insertion orientation as well as vector backbone integration. However, some large deletions are not capable to be captured by PEM-seq due to loss of primer binding sites. Additionally, rAAVs have been proved to be inefficient to transfect nonparenchymal cells accounting for nearly half of hepatocytes ^39, 40^. As we used total liver tissues to quantify targeted integration, hence the true frequencies of gene editing were likely underestimated.

In conclusion, our results demonstrate that it is a safe and effective gene therapy strategy to treat hemophilia B through integration of h*F9*p cDNAs into the r*Alb* intron 13 in rat hepatocytes via Cas9 mediated HITI strategy which also has the potential to cross-correct other genetic diseases currently treated through enzyme replacement therapeutics. Moreover, this strategy would be also critical to treat diseases which demand efficient site-specific gene integration in non-dividing cells.

## Methods

### Plasmid Construction and rAAV Vector Production

The plasmid PX458 (pSpCas9(BB)-2A-GFP, Addgene plasmid #48138) encoding SpCas9 was a gift from Feng Zhang. Six 20-nt target sequences preceding a 5’-NGG protospacer-adjacent motif (PAM) sequence located in the intron 13 of r*Alb* gene. The protospacer sequence was synthesized from Sunnybio (Shanghai, China) and inserted into PX458 through Thermo Scientific BpiI (BbsI). The sgRNA sequences and primers are shown in Supplementary Table 5. rAAV8 vectors containing the SpCas9 and either sgRNA, CMV promoter and EGFP, or the donor template, were produced as previously described using a PEI (polyethylenimine, Polysciences, 24765-2) transfection protocol followed by iodixanol gradient purification. Vectors were titrated by quantitative real-time PCR as described^41^, and the primers to measure the genome titer of rAAV8 vectors are listed in Supplementary Table 6.

### Cell Culture and Transfection

HEK293T cell lines (ATCC CRL-3216) were maintained in Dulbecco’s Modified Eagle Media (DMEM, Gibco) supplemented with 10% fetal bovine serum (FBS, Gibco), 100 U/mL penicillin, and 100 mg/mL streptomycin at 37 □ and 5% CO_2_. rAAV8 capsid (8 μg), AAV helper (12 μg), and AAV vector (10 μg) in proportion were co-transfected into HEK293T at 80% confluency in 100 mm dishes (Corning, 430167) using PEI following the manufacturer’s recommended protocol. PC12 cell lines were maintained in Roswell Park Memorial Institute (RPMI) 1640 Medium (Thermo-Gibco, 11835030) supplemented with 10% FBS, 100 U/mL penicillin, and 100 mg/mL streptomycin at 37°C and 5% CO_2_. Cells were seeded into 6-well plates (Corning, 3516) and transfected at 80% confluence using PEI. For PX458-sgRNA transfection, 800 ng of PX458-sgRNA plasmid was used. For HITI transfection, 400 ng of PX458-sgRNA and 400 ng of rAAV8-HITI-donor plasmids were cotransfected. Transfected cells were performed by Fluorescence-activated cell sorting (FACS) 72 h after transfection to enrich transfected cells.

### Animal Experiments

Sprague-Dawley (SD) strain rats purchased from Shanghai Laboratory Animal Center were housed in standard cages in a specific pathogen-free facility on a 12-hour light–dark cycle with sufficient food (irradiated) and water (autoclaved). As previously reported^42^, *F9^Δ130/Δ130^* rats were generated via CRISPR/Cas9 system using the sgRNA targeting rat *F9* gene. For systematic rAAV delivery, 8-week-old male rats received a tail vein injection of rAAV vectors at a volume of 800 μL per rat. Plasma samples for hFIX assays were collected from the orbital vein 4 weeks after rAAV injection, and every 4 weeks afterward. As previously described^43^, a two-thirds partial hepatectomy was performed on a subset of *F9^Δ130/Δ130^* rats 8 weeks after rAAV injection and liver tissues were harvested as samples for following analysis. Rats were sacrificed 36 weeks post-injection. All rat experiments conformed to the regulations drafted by the Association for Assessment and Accreditation of Laboratory Animal Care in Shanghai and were approved by the East China Normal University Center for Animal Research.

### hFIX Antigen and Activity

Plasma hFIX levels were quantified by an VisuLize™ Factor IX Antigen Kit (FIX-AG; Affinity Biologicals) and are shown as a percentage of normal level according to the manufacturer’s protocol. hFIX activity was measured by aPTT assay using a coagulation analyzer (BJ MDC, MC-4000) according to the manufacturer’s protocol. 30 μL plasma sample was mixed with 30 μL APTT reagent (BJ MDC, 03209MC) followed by a 3-min incubation at 37°C. The reaction was initiated after the addition of 30 μL of 25 mM calcium chloride (BJ MDC, 03311). Time of clot formation was recorded by the coagulation analyzer. Human plasma was used as a calibration sample.

### qPCR

Liver tissue was homogenized in liquid nitrogen and the total RNA was isolated using RNAiso Plus (Takara, catalog no. 9109). cDNA produced from a Hifair™ □ 1st Strand cDNA Synthesis Kit (YEASEN, Shanghai, China) served as a template for qPCR assays (YEASEN, Shanghai, China) to measure rat *F9, IL6, IL10, TNF-β1,* h*F9* and r*Alb*-T2A-h*F9* mRNA. Data were normalized to *β-actin* mRNA levels. All primers for qPCR are listed in Supplementary Table 6.

### Western Blot, IF, IHC, and Histology

Western blot analyses were performed on liver lysates as described previously^41^. hFIX protein was detected by a goat-anti-hFIX antibody (1:2000; Affinity Biologicals, GAFIX-AP). Mouse-anti-β-actin antibody (1:5000; Sigma, A544) was used to detect β-actin protein. For IF, liver tissues were fixed in 4% paraformaldehyde (PFA, Sigma), dehydrated in increasing concentrations of sucrose and embedded in OCT (Leica, 14020108926). 5 μm-thick serial sections were collected on slides using a cryo-sectioning machine (Danaher-Leica, CM1950). Liver slides were incubated with goat-anti-hFIX (1:2000; Affinity Biologicals, GAFIX-AP) antibody overnight at 4°C, and then incubated with FITC-anti-goat IgG secondary antibody (1:100; Boster, BA1110) for 2 h at room temperature. Slides mounted with 4’,6-diamidino-2-phenylindole (DAPI, Sigma, D9542) were imaged under the fluorescent microscope (OLYPUS, BX53F). For IHC, liver tissues were fixed in 4% paraformaldehyde (PFA, Sigma), embedded in paraplast (Leica Biosystems, 39601095) and sectioned at 5 μm. Liver slides were stained with goat-anti-hFIX antibody (1:2000, Affinity Biologicals, GAFIX-AP). Subsequently, the slides were rinsed and incubated with Biotin conjugated AffiniPure rabbit anti-goat IgG antibody (1:200; BOSTER, BA1006) and 3, 3 -diaminobenzidine substrate (Vector Labs #SK-4100, Burlingame, CA) according to manufacturer recommendations. For HE staining, liver slides were stained with hematoxylin (Solarbio, G1140) and eosin (Solarbio, G1100) according to standard protocols, and finally sealed using resin. Sections were analyzed for any abnormalities compared to rat livers from untreated group.

### Droplet digital polymerase chain reaction (ddPCR)

The general protocol of ddPCR is described according to previous report^44^. gDNAs from PC12 cells and rat livers were extracted using the TIANamp blood DNA kit, according to the manufacturer’s instructions. ddPCR reaction liquid for probes, primers, product-specific probes from Sunnybio were mixed into 35 μL reactions. Probe and primer sequences were listed in Supplementary Table 7. 30 μL oil phase mixture contained 22.5 μL oil phase A and 7.5 μL oil phase B. The ddPCR reaction liquid and the oil phase mixture were vortexed for 30 s and transiently centrifuged to remove bubbles. Loader S100 automatic sample processing system (Turtle Technology, Shanghai, China) was used to generate droplets and load samples into a chip. The condition of ddPCR reactions was as follows: 1 cycle, 50°C for 10 min, 95°C for 10 min; 45 cycles, 95°C for 20 s, 56°C for 40 s; hold at 4°C; The ramp is 2.5°C/s for every step. PCR reaction was performed using BioDigital Cycler S100 PCR amplification instrument. The proportion of inserts and targets were determined using Imager S100 Biochip Reader (Turtle Technology). Data analyses were conducted using the Imager Software version 2.2.0.1. Insertion frequency calculated by inserts/ (inserts + targets).

### PEM-seq assay

The gDNAs from the even-aged *F9^Δ130/Δ130^* rat livers without any treatment were used for the control library. 20 μg sonicated gDNAs and biotin primers (Sangon, Shanghai) were repeatedly annealed and denatured as follows: 95°C 3 min; 95°C 2 min, Ta (annealing temperature) 3 min, 5 cycles; Ta 3 min. Bst polymerase 3.0 (NEB) was added to perform for primer extension: 65°C 10 min, 80°C 5 min. AxyPrep Mag PCR Clean-Up beads (Axygen, US) were used to remove excess biotin primers. The purified products were heated to 95°C for 5 min and then rapidly cooled on ice for 5 min for DNA denaturation. Biotinylated PCR products were enriched with Dynabeads™MyOne™ Streptavidin C1 (Thermo Fisher Science, Australia). PCR products on Streptavidin C1 beads were washed with 400 μL 1 × B&W buffer (1 M NaCl, 5 mM Tris-HCl (pH=7.4), and 1 mM EDTA (pH=8.0)), then washed with 400 μL distilled water (dH_2_O), and resuspended with 42.4 μL of dH_2_O. The bridge adapter ligation reaction was progressed with 15% PEG8000 (Sigma) and T4 DNA ligase (Thermo Fisher Science, Australia) at room temperature. The ligation products were washed twice with 400 μL 1 × B&W buffer, 400 μL dH_2_O, and then resuspended with 80 μL dH_2_O. The primers of I5 and I7 sequences were used for on-beads nested PCR (Taq, Transgen Biotech, China) for 16 cycles. PCR products were then recovered using size-selection beads (Axygen, USA) and underwent tagged PCR (Fastpfu, Transgen Biotech, China) with Illumina P5 and P7 sequences. The sequences of primers were listed in Supplementary Table 8. 2 × 150 bp HiSeq was used to sequence all the PEM-seq libraries. Hiseq reads were processed according to previously reported method^26^.

### Analysis of on-target and off-target indels

PC12 cell or rat liver gDNAs was isolated using the TIANamp blood DNA kit (TIANGEN Biotech, DP304-02) following the manufacturer’s instructions. For *in vitro* on-target validation, gDNA targets were amplified through PCR for Sanger sequencing, and ICE analysis was used to analyze CRISPR gene editing efficiency as previously described^45^. For *in vivo* on-target and off-target validation, on-target and off-target sites predicted by Benchling were amplified through nested PCR, which were performed as described in Hi-TOM kits (Novogene) for targeted tagged Sequencing. The sequences of primers were listed in Supplementary Table 2. The mixture of PCR products was sequenced on the Illumina HiSeq platform as previously described^46^. All predicted potential off-target sites (OT1-11) and primer sequences used to detect indels were listed in Supplementary Table 3, 4, respectively.

### Serum biochemical analysis

For serum isolation, 1 mL whole blood was collected from the orbital vein per rat. Samples were centrifuged at 12000 rpm for 10 min at 4°C for serum collecting. The serum samples were measured for ALB, AST and ALT levels (Shanghai ADICON Clinical Laboratory, China).

### Tail-clip challenge

After anesthesia, rats were performed for a tail-clip assay as described previously^47, 48^. Briefly, the distal part of the rat tail at 1.5 mm diameter was cut and allowed to record the bleeding volume. The blood sample within 6 min was collected, and the total blood volume was measured. After holding firm pressure on the tail for 1 min, the survival rate of rats was monitored within 48 hours after clipping.

### Statistical analyses

GraphPad Prism 8.0 was used for all statistical analyses. The mean ± standard deviation (SD) was determined for each group. We used the two-tailed unpaired t-test to determine the significances between the treatment and control group. In all tests, P-values of < 0.05 were considered to be statistically significant.

## Data Availability

Targeted amplicon sequencing data and PEM-seq data have been deposited in the NCBI Sequence Read Archive database under accession codes PRJNA198802 and PRJNA707234. Sequencing data associated figures: Fig.4 b-c; Fig.5 a-b; Supplementary Fig.3 b-c; Supplementary Fig.7; Supplementary Fig. 8 b; Supplementary Fig.9 b-c. There are no restrictions on data availability. Additional information and materials will be made available upon request.

## Acknowledgements

We thank Meizhen Liu from East China Normal University for microinjection and generation of the hemophilia B rat model with support of ECNU Public Platform for innovation (011). Yin Zhang provided technical support for flow cytometry. This work was partially supported by grants from the National Key R&D Program of China (2019YFA0110802), the National Natural Science Foundation of China (No. 32025023 and 31771485), grants from the Shanghai Municipal Commission for Science and Technology (18411953500 and 20140900200).

## Author contributions

D.L., J.H., X.C., M.L., X.Z., Y.L. designed the experiments; X.C., X.N., Y.L., R.Z., L.W., L.Y., J.L., X.M., M.L, performed the experiments and analyzed the data. D.L., X.C., X.N., L.W. wrote the manuscript. D.L. supervised the research.

