## Supplementary Table_1-8 for "Amelioration of hemophilia B through CRISPR/Cas9 induced homology-independent targeted integration"

**Supplementary Table 1.** sgRNA sequences for generation of *F9*$^{\Delta 130/\Delta 130}$ rat.

| gRNA ID | Score | Location of gRNA | Target Sequence（5′-3′） | Protospacer Adaptor Motif |
|---|---|---|---|---|
| gRNA-rF9 1 | 59.9 | chrX:+143131648 | CGTCCAAAGAGATATAACTC | AGG |
| gRNA-rF9 2 | 64 | chrX:-143131783 | ATATTAATCAGGTATGATGT | GGG |

| Primers for genotyping in rats | | Length (bp) |
|---|---|---|
| ID | Sequence（5′-3′） | |
| r*F9*-KO-ID -F | TCATTATTGACACTCAACCGAT | 747 |
| r*F9*-KO-ID -R | GATGTGGACTCCTTTCCTTTT | |

**Supplementary Table 2.** Primer sequences for targeted integration analysis in the intron 13 of rAlb gene.

| **ID** | **Sequence (5'-3')** | **Length (bp)** |
|---|---|---|
| **Primers for genotyping in rats** | | |
| P-L-F | TGTGACGCATACGAACATGG | 527 |
| P-L-R | CTCGTGCTTCTTCAAAACTACACT | |
| P-R-F | GAAGATGCCAAACCAGGTCAAT | 1300 |
| P-R-R | TGGCTCTTGGCAAATGCTTATCG | |
| Hitom-*Alb-F9*p-KI-5-F | GGAGTGAGTACGGTGTGCTGCCCTGAAATGCTACTTTTTGAA | 188 |
| Hitom-*Alb-F9*p-KI-5-R | GAGTTGGATGCTGGATGGCAAGGTTTGGCCCTCCGC | |
| Hitom-*Alb-F9*p-KI-3-F | GGAGTGAGTACGGTGTGCCATTGTCTGAGTAGGTGTCATTCT | 190 |
| Hitom-*Alb-F9*p-KI-3-R | GAGTTGGATGCTGGATGGTCCATTGGAATTCTTAAAACAGGCT | |

**Supplementary Table 3.** Potential off-target sites of sgAlb7 in rat genome

| Match Name | Score | Spacer + PAM | coordinate | # of mismatch |
|---|---|---|---|---|
| sgAlb7 | 100 | GAATCATTTCACATTCCCTC CGG | chr14: 19084511 | 0 |
| *F9*_off_1 | 1.531335456 | AAGTCATTTCACATTCTCTC AGG | chr13: 99525284 | 3 |
| *F9*_off_2 | 1.445401982 | TCACCATTTTACATTCCCTC CAG | chr17: 16290752 | 4 |
| *F9*_off_3 | 1.240720994 | GAAACATTTTACATTCACTC AGG | chr15: 81461028 | 3 |
| *F9*_off_4 | 1.240350575 | GAATCATCTGTCATTCCCTC AGG | chr9: 101189192 | 3 |
| *F9*_off_5 | 1.19437619 | CAATCATTTGACCTTCCCTC AGG | chr4: 242284766 | 3 |
| *F9*_off_6 | 1.001311728 | CTATGAATTCACATTCCCTC TGG | chr9: 97607931 | 4 |
| *F9*_off_7 | 0.94878 | GATTCTTTTCCCATTCCCTC CAG | chr11:1531776 | 3 |
| *F9*_off_8 | 0.901804393 | CAACCCTCTCACATTCCCTC AGG | chr11: 36751925 | 4 |
| *F9*_off_9 | 0.894654977 | CTAACATTTCCCATTCCCTC TAG | chr18: 58108152 | 4 |
| *F9*_off_10 | 0.859852881 | GATTCATTTCAGATTCCCTA AGG | chr1: 167588939 | 3 |
| *F9*_off_11 | 0.858813601 | GTGTAATTTCTCATTCCCTC CAG | chr20: 36749557 | 4 |

**Supplementary Table 4.** Primers used for deep sequencing of on-target and potential off-target sites

| Primers for off target sites analysis in rats | |
|---|---|
| **ID** | **Sequence (5'-3')** |
| sgAlb7-OT1-F | GGAGTGAGTACGGTGTGC CCAGTCCAGGTGCTGCTAAAT |
| sgAlb7-OT1-R | GAGTTGGATGCTGGATGG ACACTTAAGCCCTCGGATGC |
| sgAlb7-OT2-F | GGAGTGAGTACGGTGTGC GCTAGGCAAGTGCTCTAC |
| sgAlb7-OT2-R | GAGTTGGATGCTGGATGG AAGGAAAATTGAAGTCACAA |
| sgAlb7-OT3-F | GGAGTGAGTACGGTGTGC TATCTGAACAATCACCTT |
| sgAlb7-OT3-R | GAGTTGGATGCTGGATGG CCTGAAAGTAGTCTGTAA |
| sgAlb7-OT4-F | GGAGTGAGTACGGTGTGC TGAGGAACCACAGTAATTAAGCCA |
| sgAlb7-OT4-R | GAGTTGGATGCTGGATGG CAGCGTTGAACACGTCCTTT |
| sgAlb7-OT5-F | GGAGTGAGTACGGTGTGC TCTGCAGGTTTGTAAGGGGAA |
| sgAlb7-OT5-R | GAGTTGGATGCTGGATGG CAGCCTGTCCTTGGATTTGC |
| sgAlb7-OT6-F | GGAGTGAGTACGGTGTGC ACAGAGTAGCTGTGAGGTCAA |
| sgAlb7-OT6-R | GAGTTGGATGCTGGATGG GTCTCCAGGCAGTTCTACCG |
| sgAlb7-OT7-F | GGAGTGAGTACGGTGTGC CCAAGGCTCAGTGAAATACTACAT |
| sgAlb7-OT7-R | GAGTTGGATGCTGGATGG AAATGTGGAATGCCTGTTATTCA |
| sgAlb7-OT8-F | GGAGTGAGTACGGTGTGC GTGCACTGGAATATCCCCAGAA |
| sgAlb7-OT8-R | GAGTTGGATGCTGGATGG CCCATGGGCCTGCTTGTTTA |
| sgAlb7-OT9-F | GGAGTGAGTACGGTGTGC ACGTCATTTGCTGGGCTTAGA |
| sgAlb7-OT9-R | GAGTTGGATGCTGGATGG TCAGACCCCTACTTCTTGTACCT |
| sgAlb7-OT10-F | GGAGTGAGTACGGTGTGC CCGCCTGTCTAGTCCACTGT |
| sgAlb7-OT10-R | GAGTTGGATGCTGGATGG GTTGTCAAGCACCGCTTTCC |
| sgAlb7-OT11-F | GGAGTGAGTACGGTGTGC TCAGGAACAGGTGTGAGTCAA |
| sgAlb7-OT11-R | GAGTTGGATGCTGGATGG TGATGGTAGTGGTGACAGCC |

**Supplementary Table 5.** The target sequences of sgRNAs and primer sequences for sequencing and T7E1 assay

| gRNA ID | Score | Location of gRNA | Target Sequence（5′-3′） | Protospacer Adaptor Motif |
|---|---|---|---|---|
| sgAlb4 | 28.47 | chr14: +19177821 | GTTCAAAAAGTAGCATTTCA | GGG |
| sgAlb6 | 24.55 | chr14: +19177765 | GAAATGATTCAGACATTGAA | TGG |
| sgAlb7 | 36.69 | chr14: -19177741 | GAATCATTTCACATTCCCTC | CGG |
| sgAlb10 | 41.79 | chr14: +19177733 | ATAAACTGTTGTTAGGGCAC | CGG |
| sgAlb11 | 34.68 | chr14: +19177727 | AAAAGGATAAACTGTTGTTA | GGG |
| sgAlb15 | 32.48 | chr14: -19177667 | GCCTGTTTTAAGAATTCCAA | TGG |

| **Primers for genotyping in rats** | | **Length (bp)** |
|---|---|---|
| **ID** | **Sequence（5′-3′）** | |
| r*Alb*-T7E1-F | TTGATTTGTGAGCCTTCC | 487 |
| r*Alb*-T7E1-R | AGCCATCGGTGATGATAC | |

**Supplementary Table 6.** Primers for qPCR and qRT-PCR.

| ID | Sequence (5′-3′) | Length (bp) |
|---|---|---|
| AAV-h*F9*p-qPCR-F | AAAATGGACTATCATATGCTTACCG | 175 |
| AAV-h*F9*p-qPCR-R | GACTCGGTGCCACTTTTTCAA | |
| AAV-Cas9-qPCR-F | GGCGTGGTTTAGGTAGTGTG | 229 |
| AAV-Cas9-qPCR-R | GTCCTCGTCCGTATTTAAGC | |
| r*β-actin*-qPCR-F | TGCTATGTTGCCCTAGACTTCG | 200 |
| r*β-actin*-qPCR-R | AGCATTGGTCACCTTTAGATGGA | |
| r*IL6*-RT-PCR-F | GTCTTCTGGAGTTCCGTTTC | 224 |
| r*IL6*-RT-PCR-R | GATGGTCTTGGTCCTTAGCC | |
| r*IL10*-RT-PCR-F | CAGGACTTTAAGGGTTACTTGG | 220 |
| r*IL10*-RT-PCR-R | CATTCTTCACCTGCTCCACT | |
| r*IFN-β*1-RT-PCR-F | TGCCATTCAAGTGATGCTCC | 169 |
| r*IFN-β*1-RT-PCR-R | CACCCAAGTCAATCTTTCCTCT | |
| r*Alb*-RT-PCR-F | GTGAGCGAGAAGGTCACCAA | 198 |
| r*Alb*-RT-PCR-R | TTTCACCAGCTCAGCGAGAG | |
| r*F9*-RT-PCR-F | TGCAGTTTTGAAGAAGCACGA | 161 |
| r*F9*-RT-PCR-R | CCAGCTTGGCACCAACATTC | |
| h*F9*-RT-PCR-F | TCCATGTTTAAATGGCGGCAG | 177 |
| h*F9*-RT-PCR-R | TCAGTACAGGAGCAAACCACC | |
| r*Alb*-h*F9*-RT-PCR-F | AGGCTGCCGACAAGGATAAC | 203 |
| r*Alb*-h*F9*-RT-PCR-R | GGTGATGAGGCCTGGTGATT | |
| r*β-actin*-RT-PCR-F | GTACGCCAACACAGTGCTG | 212 |
| r*β-actin*-RT-PCR-R | CGTCATACTCCTGCTTGCTG | |

**Supplementary Table 7.** The primer and probe sequences of ddPCR.

| **Primer** | **Sequence (5'-3')** | **Length（bp）** | **Tm** |
|---|---|---|---|
| P5F | AACTCATTTAAGCCTTGCCC | 166 | 56.28 |
| P5R | CTGTGGGAGGAAGAGAAGAG | | 56.65 |
| P3F | GGAAGAGAATAGCAGGCATG | 166 | 56.28 |
| P3R | CCTGGCCTTGCTTAATTACA | | 56.65 |
| **Probe** | **Sequence (5'-3')** | | **Tm** |
| Ins5 | Vic-TGTCTGAATCATTTC-MGB | | 64 |
| Ref3 | cy5-ACACAACAGATGTCAGAGAGCC-BHQ3 | | 64.8 |
| Ref52 | Fam-AGCTTTATCCTCTCTC-MGB | | 65 |
| Ins32 | AF594-AGCTCCGGGTCATTCTAACTAGT-BHQ2 | | 65.1 |

**Supplementary Table 8.** Primers used for PEM-seq analysis

| ID | Sequence | Note |
|---|---|---|
| LY069 | 5 Biotin -cttgtaaagcccagaatcgctaactc-3 | Forward Biotin-primer |
| LY069a | 5 -ACTCTTTCCCTACACGACGCTCTTCCGATCTAGCGcctgaaatgctactttttgaactg-3 | Forward Red-primer |
| LY069b | 5 -ACTCTTTCCCTACACGACGCTCTTCCGATCTGCCTcctgaaatgctactttttgaactg-3 | |
| LY069c | 5 -ACTCTTTCCCTACACGACGCTCTTCCGATCTAGGAcctgaaatgctactttttgaactg-3 | |
| LY069d | 5 -ACTCTTTCCCTACACGACGCTCTTCCGATCTTCAGcctgaaatgctactttttgaactg-3 | |
| LY070 | 5 Biotin -ggtaaatgcatgtgcctggccttgc-3 | Reverse Biotin-primer |
| LY070a | 5 -ACTCTTTCCCTACACGACGCTCTTCCGATCTAGCGcattggaattcttaaaacaggctc-3 | Reverse Red-primer |
| LY070b | 5 -ACTCTTTCCCTACACGACGCTCTTCCGATCTGCCTcattggaattcttaaaacaggctc-3 | |
| LY070c | 5 -ACTCTTTCCCTACACGACGCTCTTCCGATCTAGGAcattggaattcttaaaacaggctc-3 | |
| LY070d | 5 -ACTCTTTCCCTACACGACGCTCTTCCGATCTTCAGcattggaattcttaaaacaggctc-3 | |
| BA-up | 5 PO4 -CCACGCGTGCTCTACANNNTNNNNTNNNAGATCGGAAGAGCACACGT CTGAACTCCAGT-NH2 C7 3` | Bridge adapter |
| BA-down | 5 -TGTAGAGCACGCGTGGNNNNNN-NH2 C7 3` | |
| I7-B01 | 5 -CAGAAGACGGCATACGAGATCGTGATGTGACTGGAGTTCAGACGTGTGC-3 | I7-index primer<br>Nested PCR |
| P5-I5 | 5 -AATGATACGGCGACCACCGAGATCTACACACACTCTTTCCCTACACGACGC-3 | Tagged PCR Primers |
| P7-Tag | 5 -CAAGCAGAAGACGGCATACGAGAT-3 | |
