## Supplementary Fig_1-11 for "Amelioration of hemophilia B through CRISPR/Cas9 induced homology-independent targeted integration"

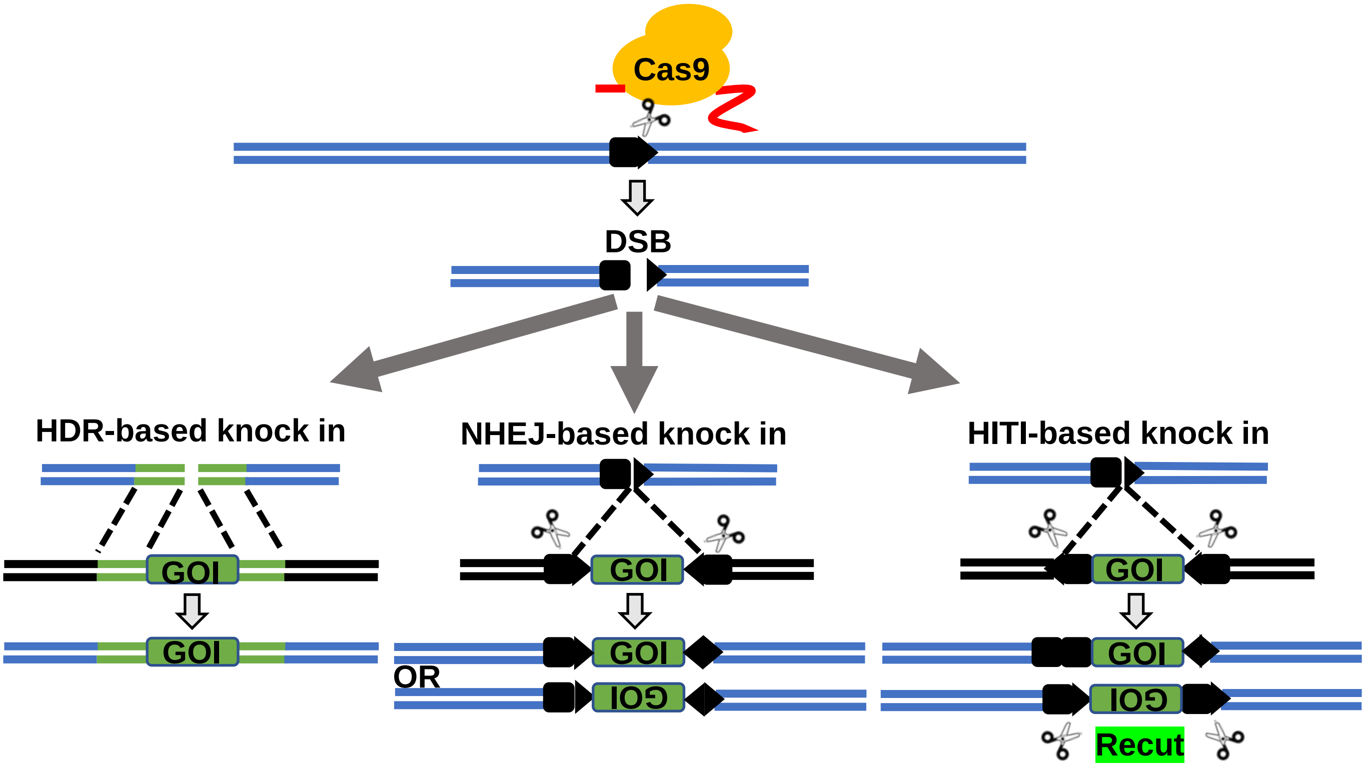


**Supplementary Fig. 1 CRISPR/Cas9-mediated targeted integration via different strategies.** CRISPR/Cas9 system induces double strand breaks (DSBs) at targeted genomic loci. These DSBs are then repaired by two major DSB repair pathways: (1) error-free HDR, which repairs these DSBs using a pair of homology arms, and (2) error-prone NHEJ, which ligates DNA ends directly. In the presence of ectopic homologous repair templates, HDR can be stimulated to insert ectopic DNA into the target site. NHEJ-mediated knock in uses a double-cut donor vector containing ectopic DNA flanked by two gRNA recognition sequences with opposite direction, resulting in the forward and/or reverse insertion of ectopic DNA at the target sites. Different with NHEJ-mediated knock in, HITI employs two sgRNA recognition sequences with the same direction and, hence, inserts ectopic DNA at target sites in the desired orientation. While the insertion of ectopic DNA in the undesired orientation would generate the intact sgRNA recognition sequence, which leads to the second round of Cas9 cutting and removes the inserted sequence. Blake pentagon, Cas9/gRNA target sequence.

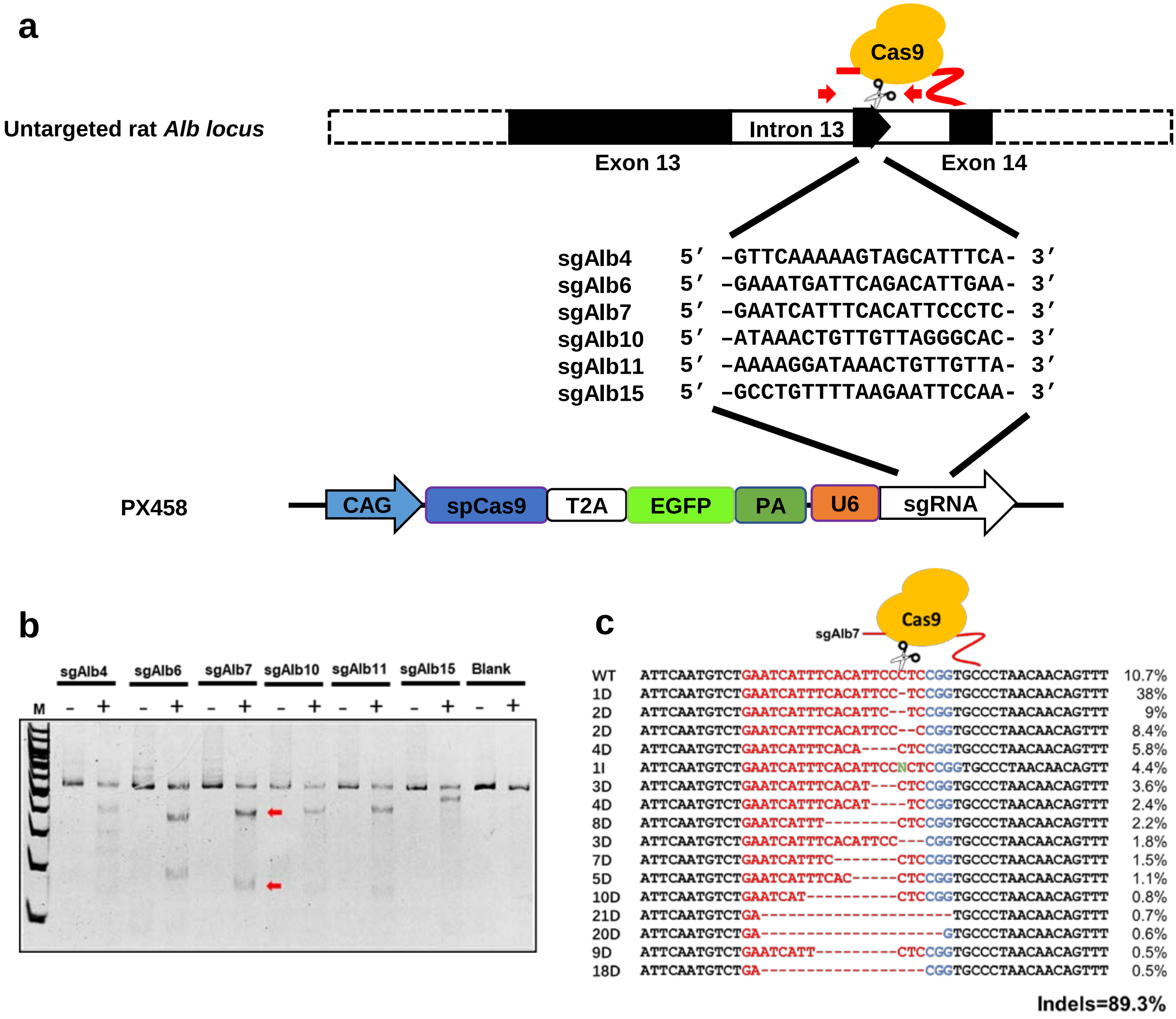


**Supplementary Fig. 2 Screening of highly efficient target sites in the intron 13 of rat *Alb* locus *in vitro*. a** CRISPR/Cas9-mediated genome editing at the site of the rat *Alb* locus. The sequences of 6 potential guide RNAs (gRNAs) were each cloned into a PX458 plasmid (Addgene plasmid #48138), and the sequences of sgRNAs were shown. Blake pentagon, Cas9/gRNA target sequence. **b** T7E1 assay was performed on rat PC12 genome obtained from samples transfected by 6 sgRNAs (sgAlb4, sgAlb6, sgAlb7, sgAlb10, sgAlb11, sgAlb15) with Cas9. **c** Sanger sequencing was analyzed for the indels frequency of sgAlb7 on rat PC12 cells by Synthego website. Arrow indicates the cleavage site of CRISPR/Cas9.

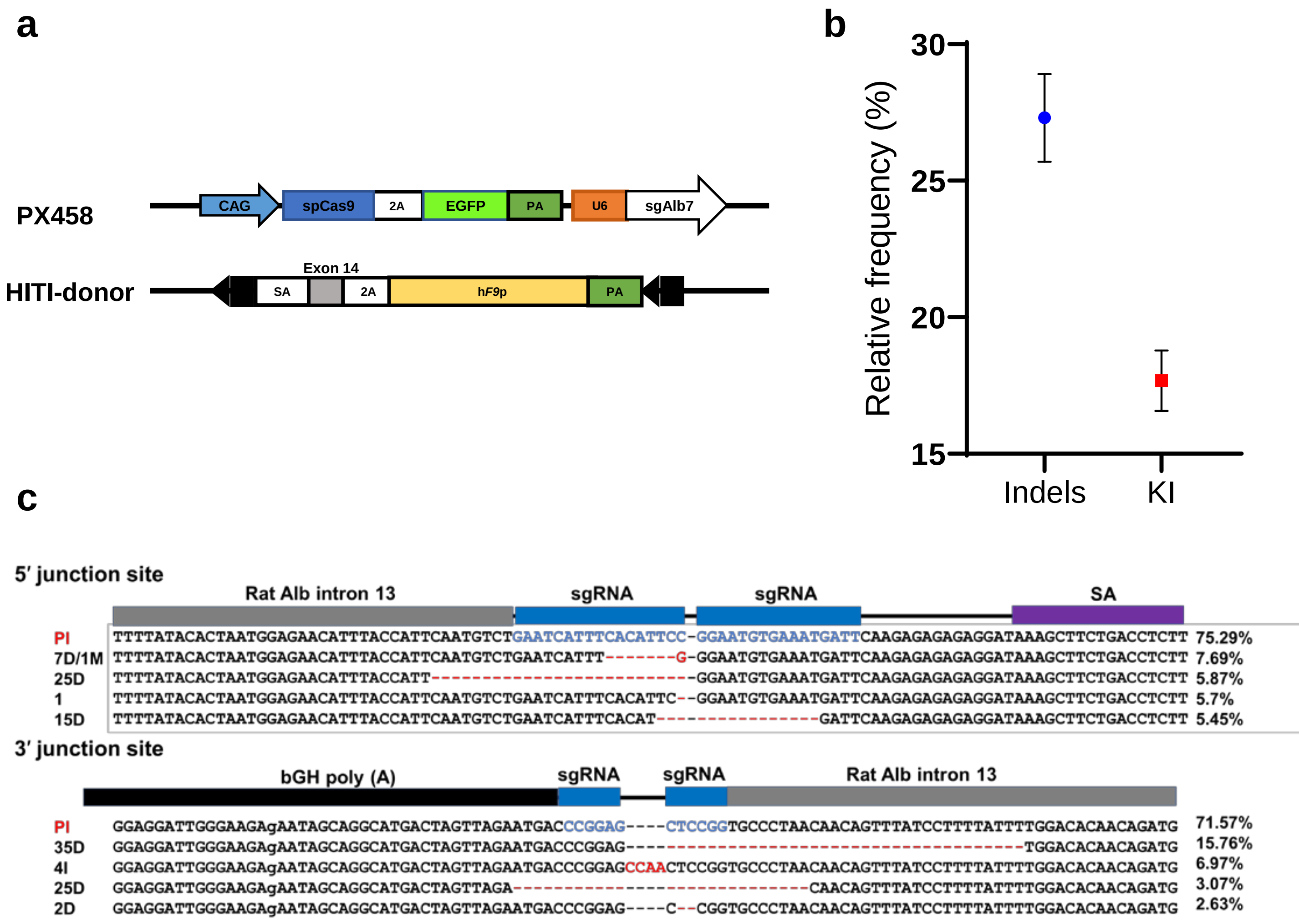


**Supplementary Fig. 3 HITI mediated knock-in of h*F9*p in rat *Alb* intron 13 *in vitro*. a** Schematic diagram of the double-cut donor and PX458 plasmids used for transfection in PC12 cells. Blake pentagon, Cas9/gRNA target sequence. **b** The efficiency of HITI-mediated knock-in measured by ddPCR assay. **c** Junction sequences of h*F9*p transgene knock-in at the intron 13 of r*Alb* locus in PC12 genome by tagged Sequencing. The Schematic diagram of HITI-mediated knocked-in is shown at the top. The partial sequence of sgRNA (17nt or 6nt) is highlighted in blue. The red lines represent the depleted sequences. Shown are representative result from one sample. PI: precise integration; D: deletion; I: insertion.

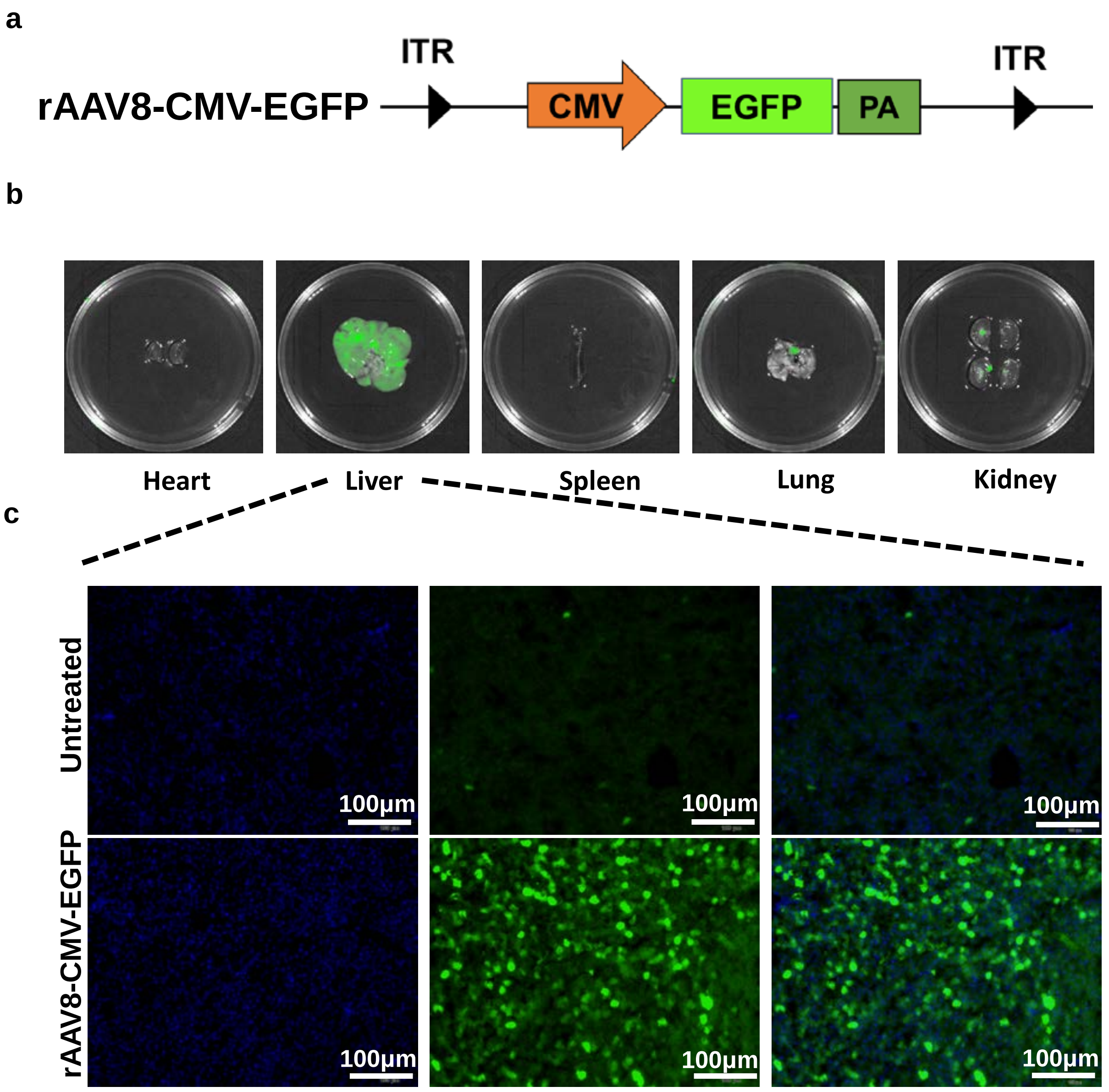


**Supplementary Fig. 4 *In vivo* infection efficiency 7 days after rAAV injection. a** Schematic of rAAV8-CMV-EGFP used for detection of infection efficiency *in vivo*. **b** Tissue imaging for eGFP in different tissues of rats by using IVIS Lumina system (PerkinElmer, IVIS Lumina LT Series II). **c** Flow cytometric analyses of eGFP in livers from untreated and rAAV-injected rats. Scale bar: 100 μm.

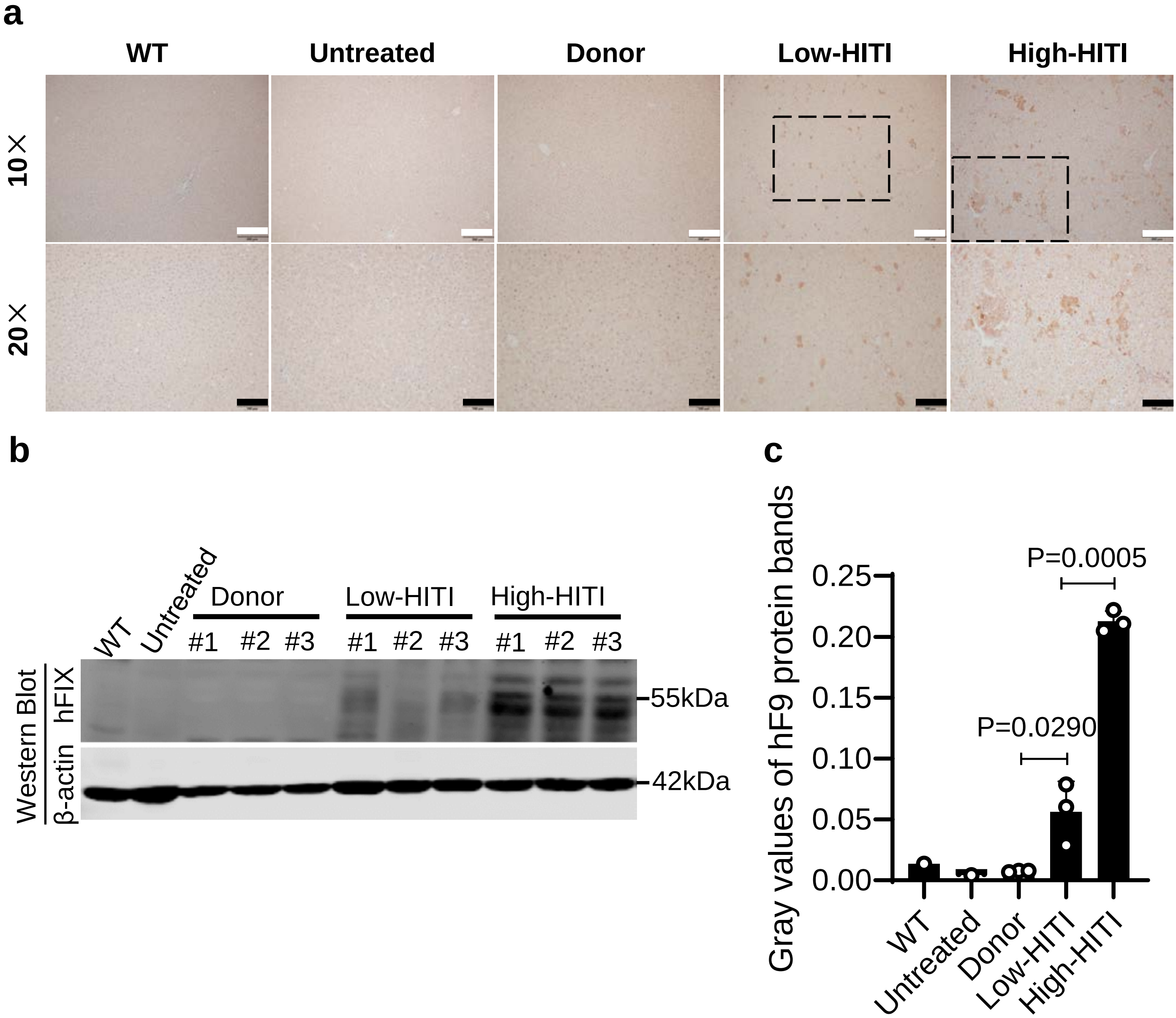


**Supplementary Fig. 5 The expression of hFIX protein in livers after rAAV injection. a** Immunohistochemistry (IHC) staining of rat liver sections from WT, untreated, donor, low-HITI and high-HITI groups, using a human-specific anti-hFIX antibody. Scale bars, 10×, 200 μm, 20×, 100 μm. **b, c** Western blot result and quantification of the hFIX protein levels in livers of the indicated groups with PHx 8 weeks after rAAV injection. hFIX protein is 55 kDa. Rat β-actin protein is used as a loading control. M, protein marker. Error bars represent SD, n=3 for donor and HITI-treated group.

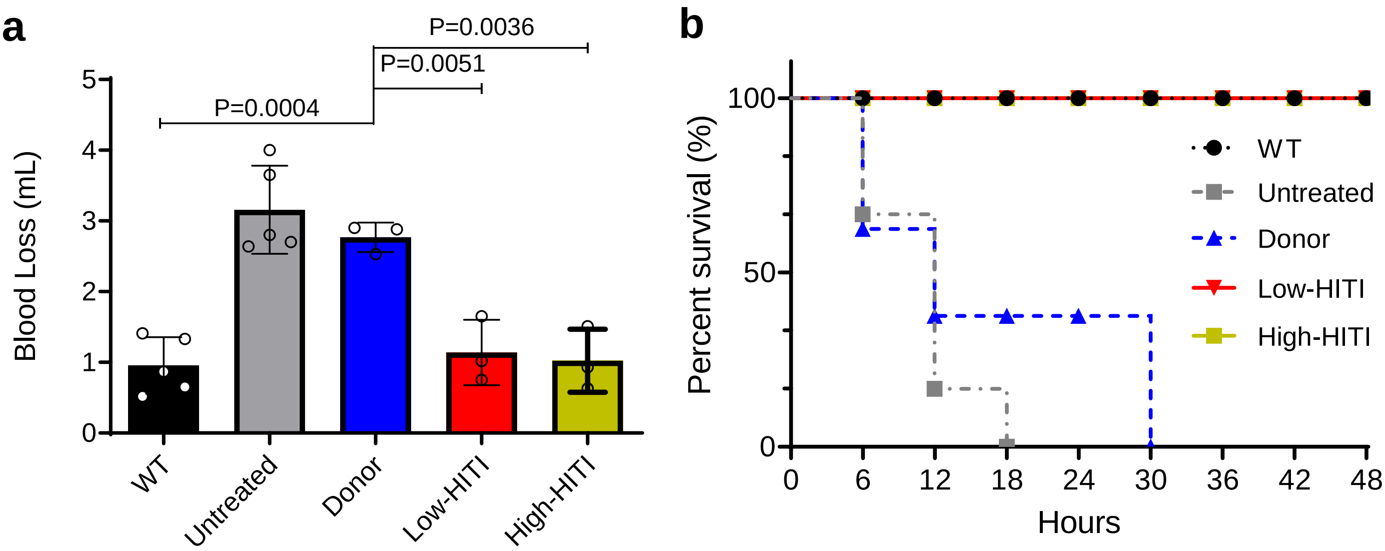


**Supplementary Fig. 6 Amelioration of coagulation function in HITI-treated rats after rAAV injection. a** Measurement of bleeding volume after tail clipping in rats 36 weeks after rAAV injection. WT, n = 5; untreated, n = 5; donor, n = 3; low-HITI, n = 3; high-HITI, n=3. Values are mean ± sd (two-tailed unpaired Student's t-test). **b** Survival rate of rats within 48 h after tail clipping 36 weeks after rAAV injection.

**a**

| | Sequence | | Reads | Indels (%) |
|---|---|---|---|---|
| Spleen | AATCATTTCACATTCC\|CTCCGG | untreated | 103205 | 0 |
| Heart | AATCATTTCACATTCC\|CTCCGG | untreated | 96720 | 0 |
| Lung | AATCATTTCACATTCC\|CTCCGG | untreated | 108343 | 0 |
| Kidney | AATCATTTCACATTCC\|CTCCGG | untreated | 134418 | 0 |
| Liver | AATCATTTCACATTCC\|CTCCGG | untreated | 146226 | 0 |

| | Sequence | | Reads | Indels (%) |
|---|---|---|---|---|
| Spleen | AATCATTTCACATTCC\|CTCCGG | HITI-high | 94149 | 0.47 |
| Heart | AATCATTTCACATTCC\|CTCCGG | HITI-high | 128397 | 1.73 |
| Kidney | AATCATTTCACATTCC\|CTCCGG | HITI-high | 106825 | 1.17 |
| Lung | AATCATTTCACATTCC\|CTCCGG | HITI-high | 114560 | 1.13 |
| Liver | AATCATTTCACATTCC\|CTCCGG | HITI-high | 111290 | 13 |

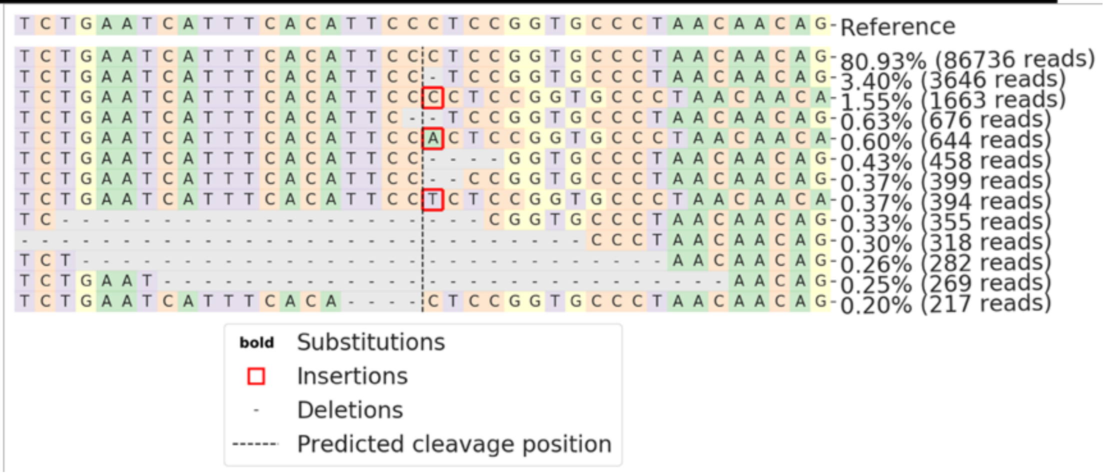


**b**

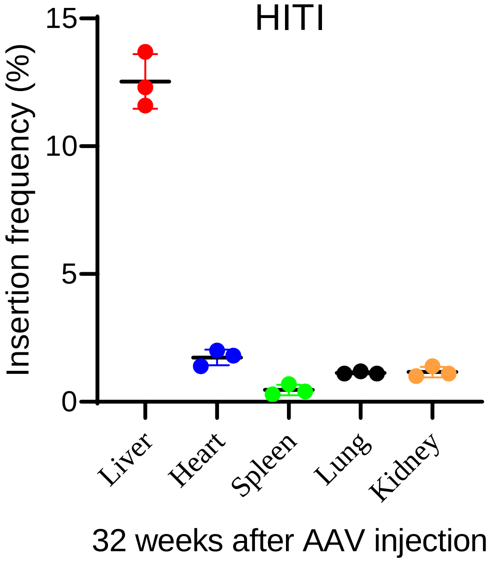


**Supplementary Fig. 7 Gene editing rarely occurs in organs other than liver after rAAV injection. a** Representative data of tagged Sequencing of the cleavage site of sgAlb7. Untreated rats served as a control. 5 major organs (liver, heart, spleen, kidney, and lung) were extracted for indel assay at 36 weeks after rAAV injection. **b** Summary of frequencies of indels induced by sgAlb7 at r*Alb* locus in 5 major organs (n = 3). Error bars represent SD.

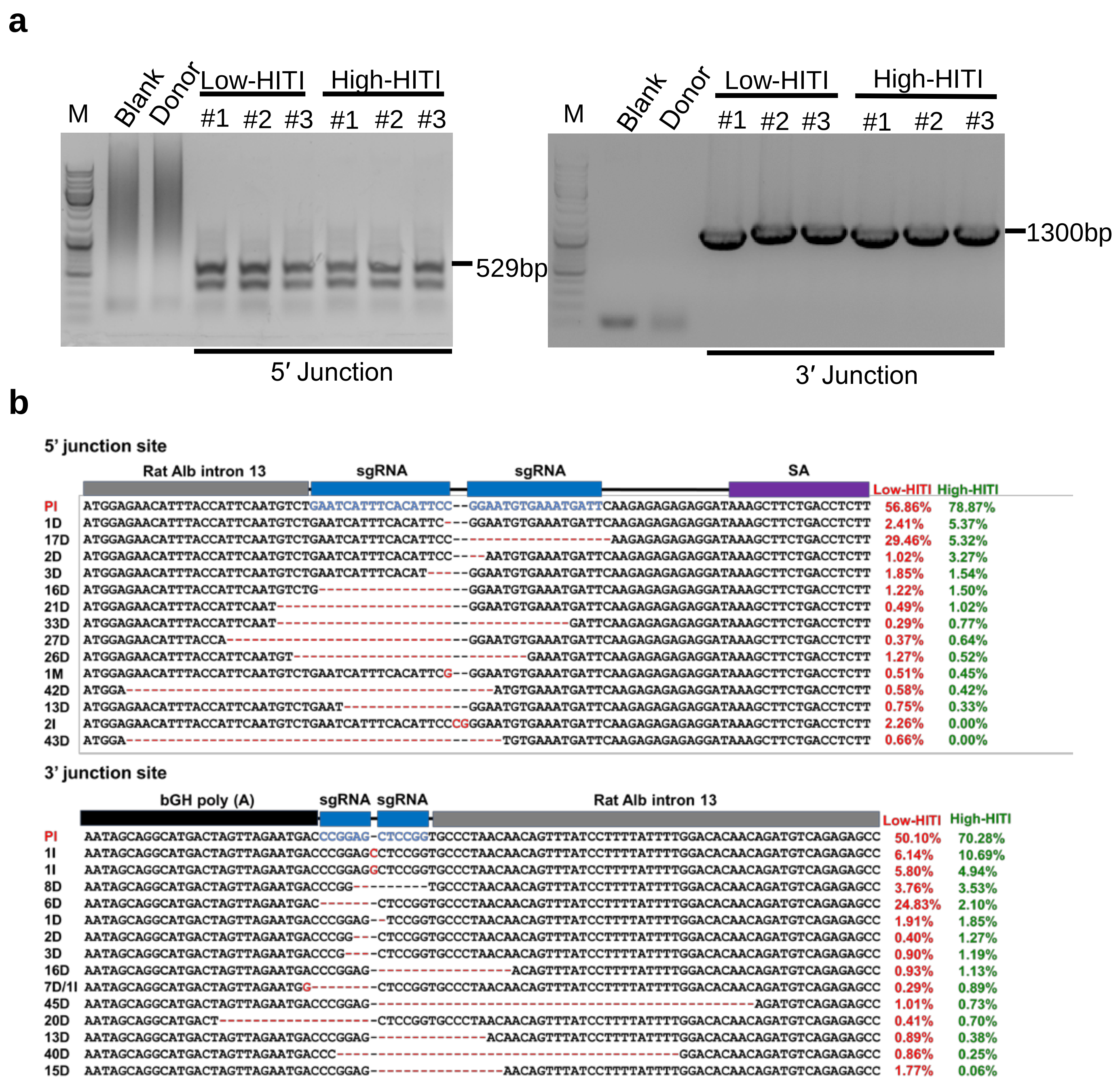


**Supplementary Fig. 8 Genotyping and Sequencing analysis of the h*F9*p knock-in via HITI at junction sites *in vivo*. a** PCR analysis with primer pairs P-L-F/R and P-R-F/P-R-R, showing successful gene targeting by HITI 8 weeks after rAAV injection at the 5′ and 3′ junction sites. Liver samples were harvested after PHx. **b** Junction sequences of h*F9*p transgene knock-in at the intron 13 of *Alb* locus in rats of HITI-treated group by tagged Sequencing. Three independent biological replicates were analyzed.

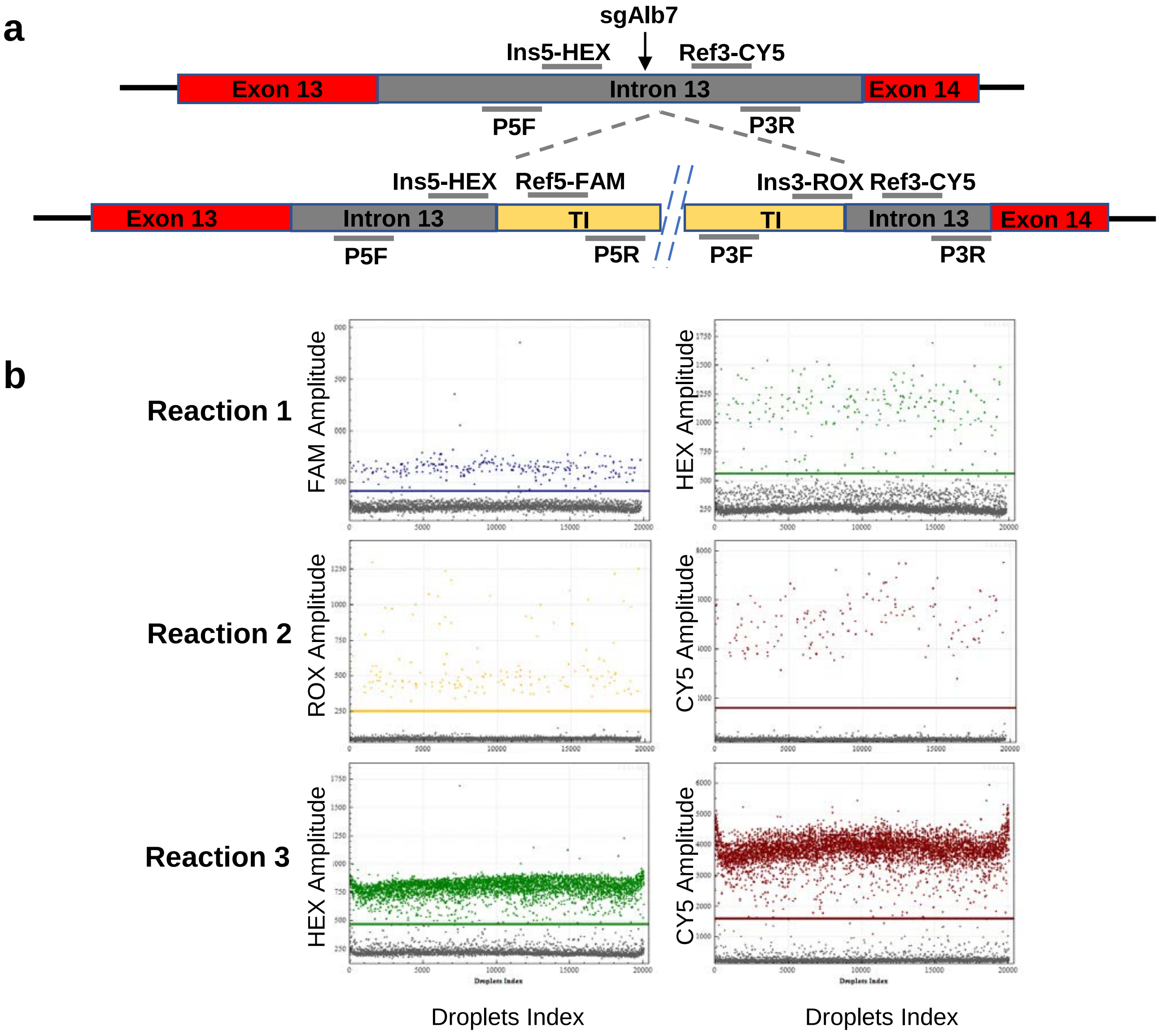


**Supplementary Fig. 9 The efficiency of targeted integration measured by ddPCR. a** Schematic illustrating the analytical strategy of targeted integration via ddPCR. Two pairs of primers and four probes were designed to amplify and detect the junctions (primers: P5F, P5R, P3F, P3R; probes: Ins5-HEX, Ref5-FAM, Ins3-ROX, Ref3-CY5). The arrow indicates the sgAlb7 target site. A representative diagram of ddPCR analysis of HITI-mediated targeted integration. Three reactions were measured in each sample: PCR products amplified from primer P5F and P5R were detected by the probe Ref5-FAM for the targeted integration copy number at 5' junction site (reaction 1, called as count-5'); PCR products amplified from primer P3F and P3R were detected by the probe Ref3-ROX for the targeted integration copy number at 3' junction site (reaction 2, called as count-3’). PCR products amplified from primer P5F and P3R were detected by the probe Ins5-HEX for the wild type copy number (reaction 3, called as count-wt); The targeted integration efficiency at the two junction sites were counted according to the following formula: 5' KI (%) = [count-5'/ (count-wt + count-5')] ×100%, and that of 3' KI (%) is the same. 100 ng of gDNA was used in each reaction. **b** A representative diagram of ddPCR analysis of the copy number of HITI-mediated knock-in.

**a**

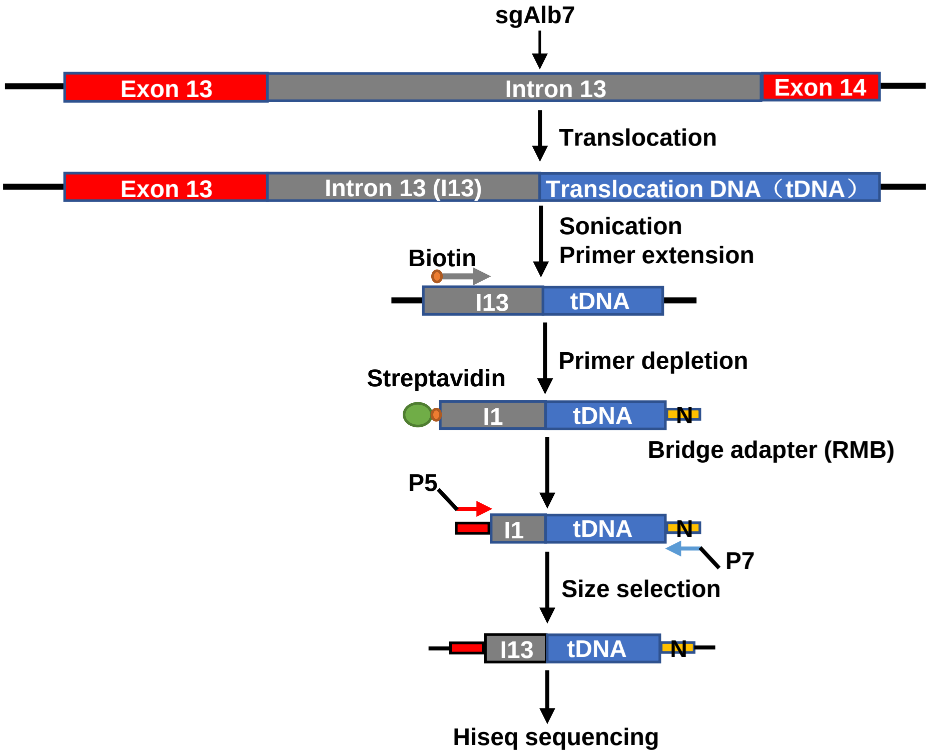


**b**

| Analytical method | Treatment | No. | Indel% | KI% | |
|---|---|---|---|---|---|
| | | | | 5' | 3' |
| PEM-seq | Low-HITI | 1 | 4.90 | 0.56 | 0.61 |
| | | 2 | 4.96 | 0.43 | 0.54 |
| | | 3 | 4.79 | 0.64 | 0.79 |
| | High-HITI | 1 | 7.91 | 0.94 | 0.96 |
| | | 2 | 10.73 | 1.66 | 1.34 |
| | | 3 | 8.08 | 0.97 | 0.98 |

**c**

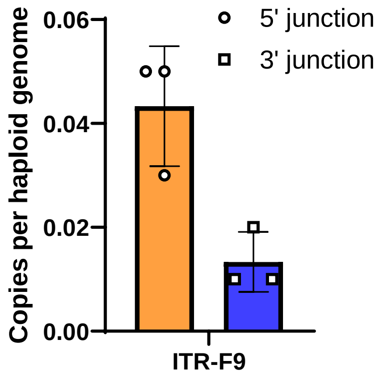


**Supplementary Fig. 10 PEM-seq assay for off-target and targeted integration. a** The Schematic diagram of PEM-seq assay. To prepare PEM-seq libraries, primer extension generated a copy of template with biotinylated primer followed by bridge adapter ligation and DNA amplification. Gray bars (intron 13) represented bait region while blue ones for captured prey region; "N" indicated random molecular barcode (RMB) in the bridge adapter. Arrows indicated positions and orientations of primers. See protocols as previously described for details. **b** The indel frequency of sgAlb7 and h*F9*p transgene knock-in efficiency (KI%) at the targeted *Alb* locus as determined by PEM-seq. **c** PEM-seq assay predicated the efficiency of h*F9*p cDNA integration mediated via AAV-ITR. Error bars represent SD, n=3 for HITI-treated group.

***hF9p* KI at intron 13 locus**

Exon13-Exon14-T2A-h*F9p*

ggctctcgctgagctggtgaaacacaagcccaaggccacagaagatcagctgaagacggtgatgggtga
cttcgcacaattcgtggacaagtgttgcaaggctgccgacaaggataactgcttcgccactgagggcc
aaaccttgttgctagaagcaaagaagccttagccgagggcagaggaagtctgctaacatgcggtgacgt
cgaggagaatcctggcccagtcgaccagcgcgtgaacatgatcatggcagaatcaccaggcctcatcac
catctgccttttaggatatctactcagtgctgaatgtacagtttttcttgatcatgaaaacgcc

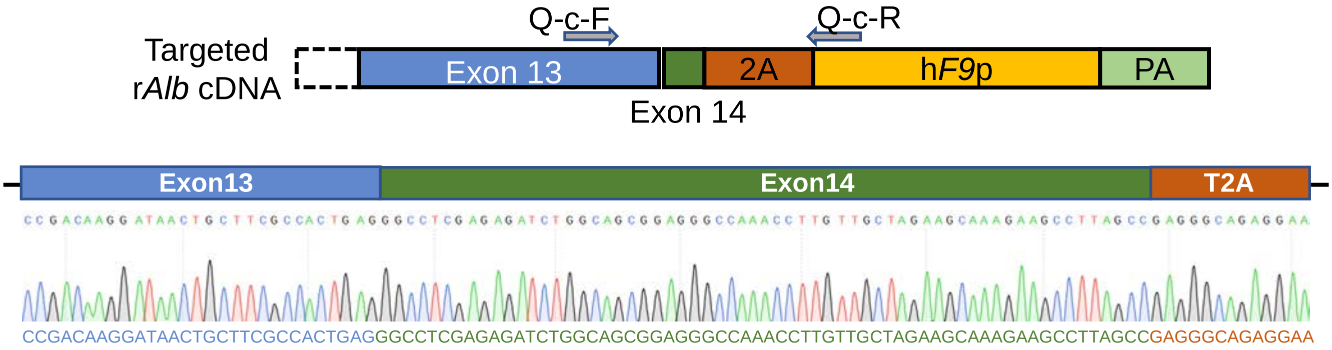


**Supplementary Fig. 11 The fusion expression of r*Alb* and h*F9*p genes was identified by RT-PCR.** Upper panel, the predicted sequence of the r*Alb*-T2A-h*F9*p fusion transcript. Lower panel, sanger sequencing demonstrated the successful transcription of r*Alb*-T2A-h*F9*p fusion transcript. Shown are representative result from one sample.
